# The developing Human Connectome Project (dHCP) automated resting-state functional processing framework for newborn infants

**DOI:** 10.1101/766030

**Authors:** Sean P. Fitzgibbon, Samuel J. Harrison, Mark Jenkinson, Luke Baxter, Emma C. Robinson, Matteo Bastiani, Jelena Bozek, Vyacheslav Karolis, Lucilio Cordero Grande, Anthony N. Price, Emer Hughes, Antonios Makropoulos, Jonathan Passerat-Palmbach, Andreas Schuh, Jianliang Gao, Seyedeh-Rezvan Farahibozorg, Jonathan O’Muircheartaigh, Judit Ciarrusta, Camilla O’Keeffe, Jakki Brandon, Tomoki Arichi, Daniel Rueckert, Joseph V. Hajnal, A. David Edwards, Stephen M. Smith, Eugene Duff, Jesper Andersson

## Abstract

The developing Human Connectome Project (dHCP) aims to create a detailed 4-dimensional connectome of early life spanning 20 to 45 weeks post-menstrual age. This is being achieved through the acquisition of multi-modal MRI data from over 1000 in- and ex-utero subjects combined with the development of optimised pre-processing pipelines. In this paper we present an automated and robust pipeline to minimally pre-process highly confounded neonatal resting-state fMRI data, robustly, with low failure rates and high quality-assurance. The pipeline has been designed to specifically address the challenges that neonatal data presents including low and variable contrast and high levels of head motion. We provide a detailed description and evaluation of the pipeline which includes integrated slice-to-volume motion correction and dynamic susceptibility distortion correction, a robust multimodal registration approach, bespoke ICA-based denoising, and an automated QC framework. We assess these components on a large cohort of dHCP subjects and demonstrate that processing refinements integrated into the pipeline provide substantial reduction in movement related distortions, resulting in significant improvements in SNR, and detection of high quality RSNs from neonates.

**Highlights:**

1. An automated and robust pipeline to minimally pre-process highly confounded neonatal fMRI data
2. Includes integrated dynamic distortion and slice-to-volume motion correction
3. A robust multimodal registration approach which includes custom neonatal templates
4. Incorporates an automated and self-reporting QC framework to quantify data quality and identify issues for further inspection
5. Data analysis of 538 infants imaged at 26-45 weeks post-menstrual age

## 1. Introduction

An increasing focus of neuroimaging science is building accurate models of the human brain’s structural and functional architecture at the macro-scale (Kaiser, 2017) through large scale neuroimaging enterprises (Van Essen et al., 2013). The mission of the developing Human Connectome Project (dHCP, http://www.developingconnectome.org) is to facilitate mapping the structural and functional development of brain systems across the perinatal period (the period before and after birth).This is being achieved through the acquisition of multi-modal MRI data from over 1000 in- and ex-utero subjects of 20–45 weeks post-menstrual age (PMA), combined with the development of optimised pre-processing pipelines. The ambitious scale of the project will enable developing detailed normative models of the perinatal connectome. The raw and processed data from the project, along with genetic, clinical and developmental information (Hughes et al., 2017), will be made publicly available via a series of data releases.

As the human infant enters the world, core functional neural systems are rapidly developing to provide essential functional capabilities. Characterisation of the perinatal brain using fMRI can provide insights into the relative developmental trajectories of brain systems during this crucial period of development (Cusack et al., 2017). FMRI has been used to characterise the neural activity associated with the sensorimotor systems (Arichi et al., 2010), olfaction (Arichi et al., 2013), and visual (Deen et al., 2017), auditory (Anderson et al., 2001), vocal (Dehaene-Lambertz et al., 2002), and emotional perception (Blasi et al., 2011; Graham et al., 2013). However, whilst task-based studies are informative, they are difficult to perform in young, pre-verbal infants. Studies of spontaneous brain activity are ideally suited to the perinatal period and can provide an overall view of the spatial and temporal organisation of functional systems and their maturation. Using this approach, a number of studies have explored the emergence of the resting-state functional networks (RSNs) in infants (Fransson et al., 2007; Lin et al., 2008; Liu et al., 2008). These RSNs are found to be emerging in the preterm period and are largely present at the age of normal birth (40 weeks PMA), (Doria et al., 2010; Fransson et al., 2007; Gao et al., 2015; He and Parikh, 2016; Smyser et al., 2010), increasing in strength over the first year of life (Damaraju et al., 2014).

Acquisition, pre-processing, and analysing MRI data from the fetal and neonatal population presents unique challenges as the tissue composition, anatomy, and function undergo rapid changes during the perinatal period and markedly differ from those in the adult brain (Ajayi-Obe et al., 2000; Dubois et al., 2014; Gilmore et al., 2012; Inder et al., 1999; Kapellou et al., 2006). These differences demand re-evaluation of established pipelines (Cusack et al., 2017; Mongerson et al., 2017; Smyser et al., 2016). Changes in tissue composition, due to processes such as myelination, and neural and vascular pruning (Dubois et al., 2014; Kozberg and Hillman, 2016) affect imaging contrast (Goksan et al., 2017; Rivkin et al., 2004). These changes require bespoke developmental structural templates (Kuklisova-Murgasova et al., 2011; Schuh et al., 2018a; Shi et al., 2018) and optimised registration techniques (Deen et al., 2017; Goksan et al., 2015). Care is required to ensure that the effects of changing relative voxel resolution and SNR on analyses are ameliorated and monitored (Cusack et al., 2017; Gao et al., 2015). Infant brain haemodynamics differ from adults, and can show substantial changes over the perinatal period (Arichi et al., 2012; Cornelissen et al., 2013; Kozberg and Hillman, 2016). Importantly, levels of head motion over extended fMRI scans are typically high and differ in nature from adults (Cusack et al., 2017; Deen et al., 2017; Satterthwaite et al., 2012; Smyser et al., 2010). As motion and pulsatile artefacts can have profound effects on measures of resting-state connectivity, great care with motion and distortion correction is required in the neonate (Deen et al., 2017; Power et al., 2012).

A major focus of the dHCP project is therefore the advancement of acquisition and analysis protocols optimised for the infant brain (Bastiani et al., 2018; Bozek et al., 2018; Hughes et al., 2017; Makropoulos et al., 2018). The present report provides a detailed description of the dHCP resting-state functional MRI (rfMRI) pre-processing pipeline for neonates. The pipeline is inspired by the Human Connectome Project (HCP) minimal pre-processing pipelines (Glasser et al., 2013) and the FSL FEAT pipeline (Jenkinson et al., 2012) for adults; however it is designed to specifically address the challenges that neonatal data present. Each stage of the pipeline has been assessed and refined to ensure a high level of performance and reliability. The pipeline includes integrated dynamic distortion and motion correction, a robust multimodal registration approach, bespoke ICA-based denoising, and an automated QC framework. We assess these components, showing results from an initial cohort of dHCP subjects. The processed data from these pipelines are currently available for download. We apply PROFUMO (Harrison et al., 2015), a Bayesian group component decomposition algorithm (with a customised neonatal HRF prior), to demonstrate high quality RSNs from these data. A companion paper (Baxter et al., 2019) assesses the pipeline, applying it to a stimulus response dataset. In order to present the clearest description of the pipeline stages throughout the paper, we do not separate out Methods and Results sections, but intermix descriptions of methods, their assessment procedures and results.

## 2. Subjects and fMRI acquisition

### 2.1. Subjects

MR images were acquired as a part of the dHCP which was approved by the National Research Ethics Committee and informed written consent given by the parents of all participants.

Data from two cohorts of dHCP subjects are used in this paper, referred to as dHCP-538 and dHCP-40. The dHCP-538 is a large cohort that comprises 538 scans and is used to evaluate overall performance of the dHCP neonatal fMRI pipeline, as well as to assess most processing stages. The dHCP-40 is a smaller subset that comprised 40 scans and is used to specifically contrast and evaluate the more computationally demanding motion and distortion correction algorithms (see Section 3.4).

The dHCP-538 cohort comprises 538 scans that passed upstream QC prior to the dHCP neonatal fMRI pipeline described in this paper (see supplementary section 9.2), and had been processed with the pipeline as of the time of writing. These 538 scans were obtained from 422 subjects scanned once and 58 subjects scanned twice (480 subjects in total). The first scan was pre-term, <37 weeks PMA, and second scan was term equivalent age. The dHCP-538 contains 215 females and 265, and has a mean PMA at scan of 39.81 weeks (*σ*=3.36). This cohort is a superset of the 1^st^ (2017) and 2^nd^ (2019) dHCP public data releases. The dHCP-40 scans are from 40 subjects (all scanned once) that were released in the 1^st^ dHCP data release, in 2017. The dHCP-40 contains 15 females and 25 males and has a mean PMA at scan of 39.81 weeks (*σ*=2.17). The joint distribution of age-at-birth and age-at-scan for both cohorts is presented in Figure 1

**Figure 1.**
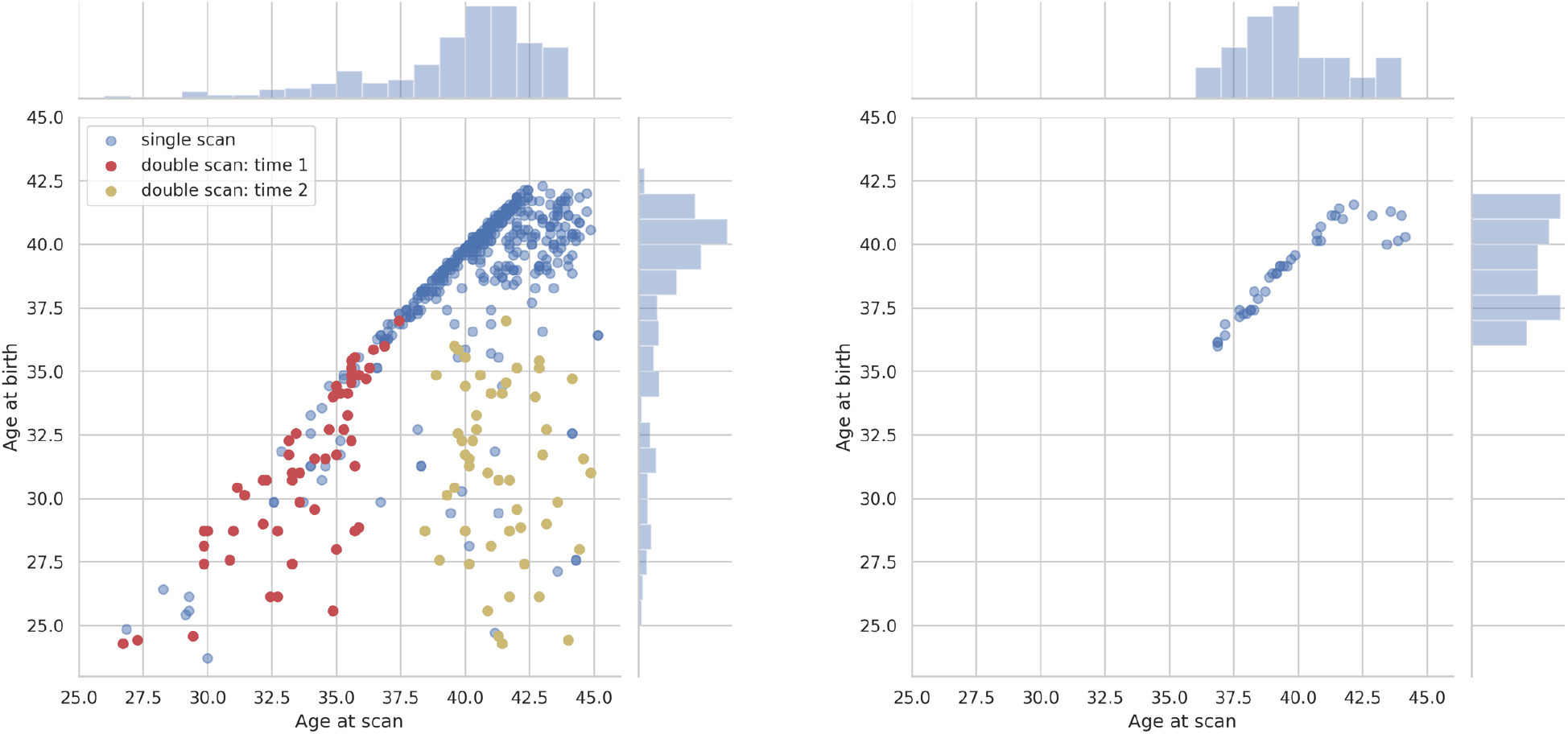
Joint distributions of post-menstrual age-at-birth (weeks) and post-menstrual age-at-scan (weeks) for the dHCP-538 (left) and dHCP-40 (right) cohorts.

### 2.2. Acquisition protocol summary

All data were acquired on a 3T Philips Achieva with a dedicated neonatal imaging system including a neonatal 32-channel phased-array head coil (Hughes et al., 2017), sited within the neonatal intensive care unit at the Evelina London Children’s Hospital. Anatomical images (T1w and T2w), resting-state functional (rfMRI) and diffusion acquisitions were acquired without sedation, with a total examination time of 63mins.

#### Anatomical acquisition and pre-processing

T2w (TR=12s; TE=156ms; SENSE factor: axial=2.11, sagittal=2.58) and inversion recovery T1w (TR=4795ms; TI=1740ms; TE=8.7ms; SENSE factor: axial=2.26, sagittal=2.66) multi-slice fast spin-echo images were each acquired in sagittal and axial slice stacks with in-plane resolution 0.8×0.8mm^2^ and 1.6mm slices overlapped by 0.8mm (see Table 1 for a summary). Both T2w and T1w images were reconstructed using a dedicated neonatal motion correction algorithm (Cordero-Grande et al., 2018). Retrospective motion-corrected reconstruction (Cordero-Grande et al., 2018) and integration of the information from both acquired orientations (Kuklisova-Murgasova et al., 2012) were used to obtain 0.8 mm isotropic T2w and T1w volumes with significantly reduced motion artefacts. Anatomical pre-processing of the T2w and T1w images was performed using the dHCP neonatal structural processing pipeline (Makropoulos et al., 2018). Several outputs of the dHCP neonatal structural pipeline are required by the dHCP neonatal fMRI pipeline described in this paper; specifically, the bias corrected T2w image in native space, the bias corrected T1w image sampled to T2w native space, and the T2w tissue segmentation (9 labels). The structural pipeline outputs used by this neonatal fMRI pipeline were pre-processed with dHCP structural pipeline version 1.1 (Makropoulos et al., 2018).

**Table 1.**
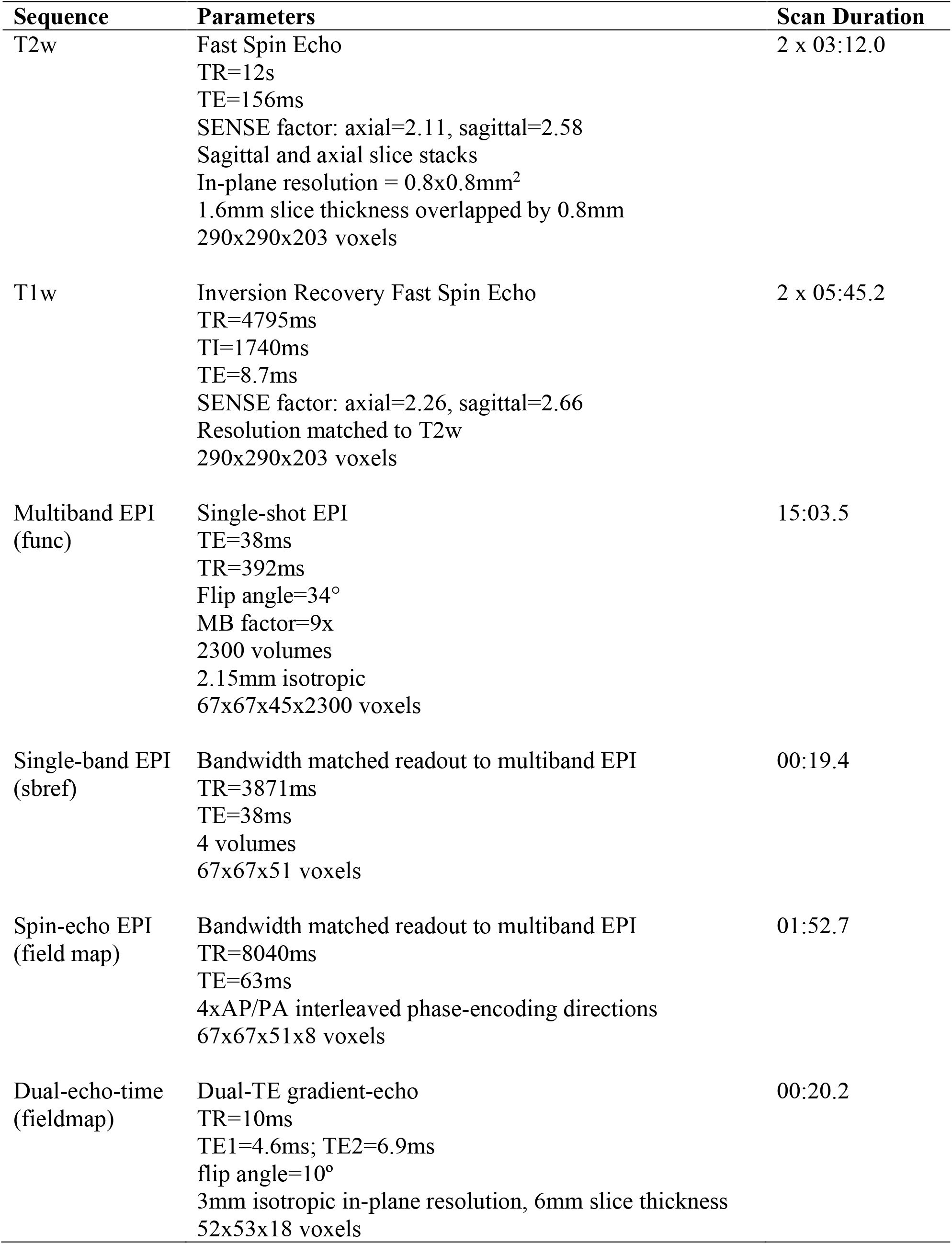
Acquisition sequence parameters

| Sequence | Parameters | Scan Duration |
| --- | --- | --- |
| T2w | Fast Spin Echo<br>TR=12s<br>TE=156ms<br>SENSE factor: axial=2.11, sagittal=2.58<br>Sagittal and axial slice stacks<br>In-plane resolution = 0.8x0.8mm<sup>2</sup><br>1.6mm slice thickness overlapped by 0.8mm<br>290x290x203 voxels | 2 x 03:12.0 |
| T1w | Inversion Recovery Fast Spin Echo<br>TR=4795ms<br>TI=1740ms<br>TE=8.7ms<br>SENSE factor: axial=2.26, sagittal=2.66<br>Resolution matched to T2w<br>290x290x203 voxels | 2 x 05:45.2 |
| Multiband EPI<br>(func) | Single-shot EPI<br>TE=38ms<br>TR=392ms<br>Flip angle=34°<br>MB factor=9x<br>2300 volumes<br>2.15mm isotropic<br>67x67x45x2300 voxels | 15:03.5 |
| Single-band EPI<br>(sbref) | Bandwidth matched readout to multiband EPI<br>TR=3871ms<br>TE=38ms<br>4 volumes<br>67x67x51 voxels | 00:19.4 |
| Spin-echo EPI<br>(field map) | Bandwidth matched readout to multiband EPI<br>TR=8040ms<br>TE=63ms<br>4xAP/PA interleaved phase-encoding directions<br>67x67x51x8 voxels | 01:52.7 |
| Dual-echo-time<br>(fieldmap) | Dual-TE gradient-echo<br>TR=10ms<br>TE1=4.6ms; TE2=6.9ms<br>flip angle=10°<br>3mm isotropic in-plane resolution, 6mm slice thickness<br>52x53x18 voxels | 00:20.2 |

#### rfMRI

High temporal resolution multiband EPI (TE=38ms; TR=392ms; MB factor=9x; 2.15mm isotropic) specifically developed for neonates (Price et al., 2015) was acquired for 15 minutes. No in-plane acceleration or partial Fourier was used. Single-band EPI reference (sbref) scans were also acquired with bandwidth-matched readout, along with additional spinecho EPI acquisitions with 4xAP and 4xPA phase-encoding directions. Reconstructions follow the extended SENSE framework (Zhu et al., 2016) with sensitivity maps computed from the matched single-band data. Field maps were obtained from an interleaved (dual TE) spoiled gradient-echo sequence (TR=10ms; TE1=4.6ms; TE2=6.9ms; flip angle=10°; 3mm isotropic in-plane resolution, 6mm slice thickness). Phase wraps were resolved by solving a Poisson’s equation (Ghiglia and Romero, 1994).

## 3. Pre-processing pipeline

### 3.1. Pipeline overview

The dHCP neonatal fMRI pipeline is inspired by the HCP minimal pre-processing pipelines (Glasser et al., 2013) and the FSL FEAT pipeline (Jenkinson et al., 2012) for adults; however it is designed to specifically address the challenges that neonatal data presents. These challenges and their solutions are detailed throughout the paper.

The goal of the pipeline is to generate high-quality minimally pre-processed rfMRI data for open-release to the neuroimaging community. The motivation for “minimal” pre-processing is to ensure that the scientific community will not be restricted in the subsequent analysis (including further pre-processing) that they can perform on the data. Therefore, in building the pipeline we have restricted ourselves to the pre-processing steps that we consider absolutely crucial for the widest possible range of subsequent analyses.

The inputs to the pipeline are the raw multi-band EPI functional (func), single-band EPI reference (sbref), and spin-echo EPI with opposing phase-encode directions, as well as the dHCP structural pipeline pre-processed outputs: bias corrected T2w structural image (struct), bias corrected T1w image aligned with the T2w, and the T2w discrete segmentation (dseg).

The primary output is the minimally pre-processed 4D functional image which is motion corrected, distortion corrected, high-pass filtered and denoised. The secondary outputs are transforms to align the pre-processed functional images with the structural (T2w) and template (atlas) spaces.

A schematic of the dHCP neonatal fMRI pipeline is presented in Figure 2. The main parts of the pipeline are:

1. Fieldmap pre-processing: estimate the susceptibility distortion field and align it with the functional data (see Section 3.2)
2. Registration: align all images with the native T2 space and the neonatal atlas space (see Section 3.3)
3. Susceptibility and motion correction: Perform slice-to-volume motion correction and dynamic susceptibility distortion correction, and estimate motion nuisance regressors (see Section 3.4)
4. Denoising: Estimate artefact nuisance regressors and regress all nuisance regressors from the functional data (see Section 3.5)

**Figure 2.**
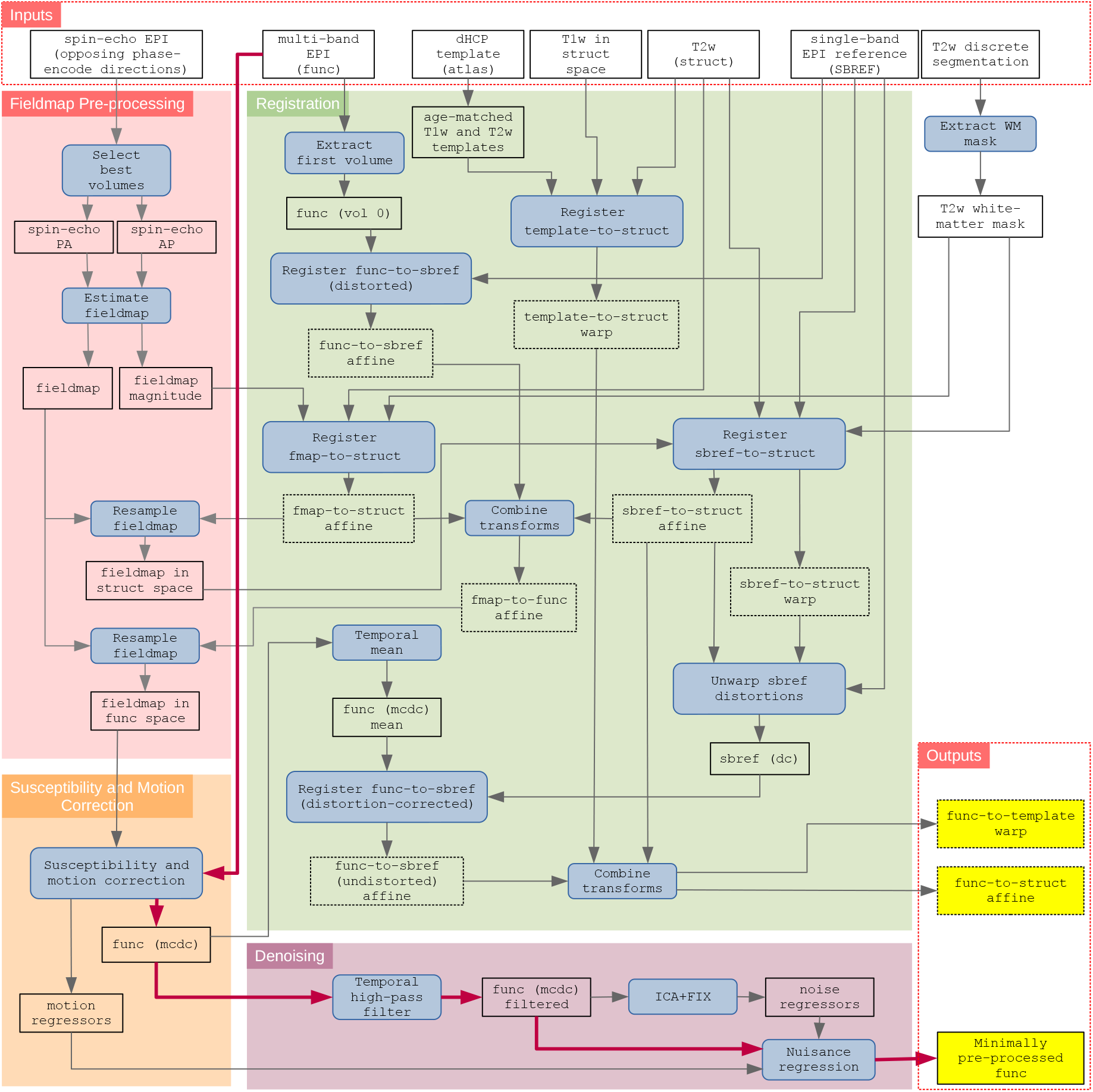
Schematic of the dHCP fMRI neonatal pre-processing pipeline. The schematic is segregated into the 4 main conceptual processing stages by coloured background; fieldmap pre-processing (red), susceptibility and motion correction (orange), registration (green), and denoising (purple). Inputs to the pipeline are grouped in the top row, and the main pipeline outputs are grouped in the lower right. Blue filled rectangles with rounded corners indicate processing steps, whilst black rectangles (with no fill) represent data. The critical path is denoted by magenta connector arrows. (dc) = distortion corrected; (mcdc) = motion and distortion corrected.

### 3.2. Fieldmap pre-processing

The EPI sequence is sensitive to field inhomogeneities caused by differences in magnetic susceptibility across the infant’s head. This results in distortions in the image in the phase-encode (PE) direction, particularly at tissue interfaces. However, if the susceptibility-induced off-resonance field is known, these distortions are predictable and can be corrected.

The dHCP neonatal fMRI pipeline uses FSL TOPUP (Andersson et al., 2003) to estimate the susceptibility-induced off-resonance field from the spin-echo EPI with opposing phase-encoding directions, and converts that to a voxel displacement field to correct the EPI distortions. The input to TOPUP is two volumes of the spin-echo EPI for each phase-encoding direction. These volumes will have different distortions because of the differing PE directions, so TOPUP uses an iterative process to estimate an off-resonance field that minimises the distortion corrected difference between the two images.

The dHCP spin-echo EPI has 8 volumes with 2 PE directions (4 x AP, 4 x PA). Movement of the subject during acquisition results in striping artefact in the PE direction (see Supplementary Figure 1). The two-best spin-echo EPI volumes (1 per PE direction) are selected as inputs to TOPUP. Here, “best” is defined as the smoothest over the z-dimension, which avoids motion artefact characterised by intensity differences between slices, a characteristic “stripy” appearance. This z-smoothness metric is obtained per volume by calculating the voxel-wise standard deviation of the slice-to-slice difference in the z-dimension and then selecting the minimum standard deviation per volume. Distributions of z-smoothness are presented in Supplementary Figure 2.

The method described above worked well for selecting the best spin-echo EPI volume pair, but it was difficult to find a threshold to determine if this pair was still “good enough”. It was therefore combined with visual inspection, and 12.7% (75 of 590) were visually identified as having significant movement contamination in all of the volumes for the given subject. In this circumstance, the fall-back procedure was to use the dual-echo-time-derived fieldmap instead of the spin-echo-EPI-derived fieldmap. Where possible, the spin-echo-EPI-derived fieldmap was used in preference to the dual-echo-time-derived fieldmap due to higher anatomical contrast in the magnitude image allowing more reliable registration to the structural T2w image (see Supplementary Figure 3). Furthermore, the lack of contrast in the dual-echo-time-derived fieldmap magnitude meant that it was often impossible to adequately judge the quality of the registration.

To ensure that the dual-echo-time and spin-echo-EPI derived fieldmaps could be used interchangeably, we evaluated the similarity between the two and found them to be qualitatively and quantitatively similar (see supplementary section 9.1).

### 3.3. Registration

There are two main target volumetric alignment spaces within the dHCP neonatal fMRI pipeline (see Table 2); 1) the within-subject structural space defined as the subject’s native T2w space, and 2) the between-subject group standard space defined as the 40-week template from the dHCP volumetric atlas (Schuh et al., 2018a). We refer to these spaces as **structural** and **template** respectively.

**Table 2.** Spaces and transforms used in the dHCP neonatal fMRI pipeline. Superscript (−1) refers to the inverse of the transform.

| Spaces |  |  |
| --- | --- | --- |
| functional (func) | Native multiband EPI space |  |
| sbref | Native single-band EPI reference space |  |
| structural (struct) | Native T2w space |  |
| fieldmap (fmap) | Derived fieldmap space |  |
| template | dHCP 40-week template space |  |
| Primary Registrations |  | Degrees of freedom |
| (a) | fieldmap-to-structural | rigid |
| (b) | sbref-to-structural | rigid |
| (c) | functional-to-sbref (distorted) | rigid |
| (d) | functional-to-sbref (undistorted) | rigid |
| (e) | template-to-structural | nonlinear |
| Composite Registrations |  |  |
| $(a) \oplus (b)^{-1} \oplus (c)^{-1}$ | fieldmap-to-functional | rigid |
| $(d) \oplus (b)$ | functional-to-structural (undistorted) | rigid |
| $(d) \oplus (b) \oplus (e)^{-1}$ | functional-to-template (undistorted) | nonlinear |

The brain is undergoing rapid developmental changes during the perinatal period, that require explicit consideration when registering to these spaces. Specifically,

1. The myelination and water content of the white matter is still maturing, resulting in inversion of T1w/T2w MRI contrast when compared to adult brain scans. The impact of this inhomogeneous myelination can be mitigated by using the T2w image as the structural target space, as opposed to the T1w which is more typical in adult cohorts. Furthermore, we use the BBR cost-function for intra-subject registrations, which only samples the image intensity along the high-contrast GM/WM boundary to create an intensity gradient, and is more resistant to the inhomogeneous myelination than other registration cost functions that uniformly sample the whole image.
2. The brain increases greatly in both size and gyrification during the perinatal period, which makes it challenging to define an unbiased common template space for group analyses. Therefore, we use a bespoke developmental atlas developed using dHCP data (Schuh et al., 2018a)(see Figure 5). The dHCP volumetric atlas contains T1w/T2w volumetric templates per week from 36-43 weeks PMA. We have augmented it with week-to-week nonlinear transforms estimated using a diffeomorphic multi-modal (T1w/T2w) registration (ANTs SyN) (Avants et al., 2008).

An additional registration challenge is that the dHCP uses a fast-multi-band EPI sequence which is advantageous with regard to minimising the impact of motion (see section 3.4), but which results in poorer tissue contrast. We mitigate this by using a single-band reference image (sbref) as an intermediate registration target, as per the adult HCP pre-processing pipelines (Glasser et al., 2013).

Another consideration when developing the registration protocol was to ensure reliability across a large cohort so that we could minimise manual intervention. We found empirically that BBR was more robust than other registration cost functions. We attribute this to the fact that BBR only samples the image along the more reliable high-contrast WM/GM boundary and is therefore less susceptible to image defects. BBR performance was evaluated with detailed visual scoring of registration quality by multiple judges on a subset of subjects.

To achieve alignment to structural and template spaces we perform five primary registrations (see Table 2): (1) fieldmap-to-structural, (2) sbref-to-structural, (3) functional-to-sbref (distorted), (4) functional-to-sbref (distortion-corrected), and (5) template-to-structural. In the event that the age of the subject is outside the range covered by the atlas, the closest template age within the atlas is used. From these five primary registrations, a variety of composite alignments can be calculated, most importantly: (1) fieldmap-to-functional, (2) functional-to-structural (undistorted), and (3) functional-to-template (undistorted). These registration steps are described schematically in the green box of Figure 2, and further detail is presented in Supplementary Section 9.3.

Registration quality was assessed on the dHCP-538 dataset for each of the primary registrations by evaluating the similarity of the source (moving) image (re-sampled to reference space) and the reference (fixed) image. Normalised mutual information (NMI) was used as a metric of similarity.

The distribution of the NMI for each of the primary registration steps is presented in Figure 3. All the distributions appear unimodal and there are very few outliers on the lower tail (i.e., less similar). The pipeline flags these outliers for manual investigation. Furthermore, these registrations were also manually visually checked.

**Figure 3.**
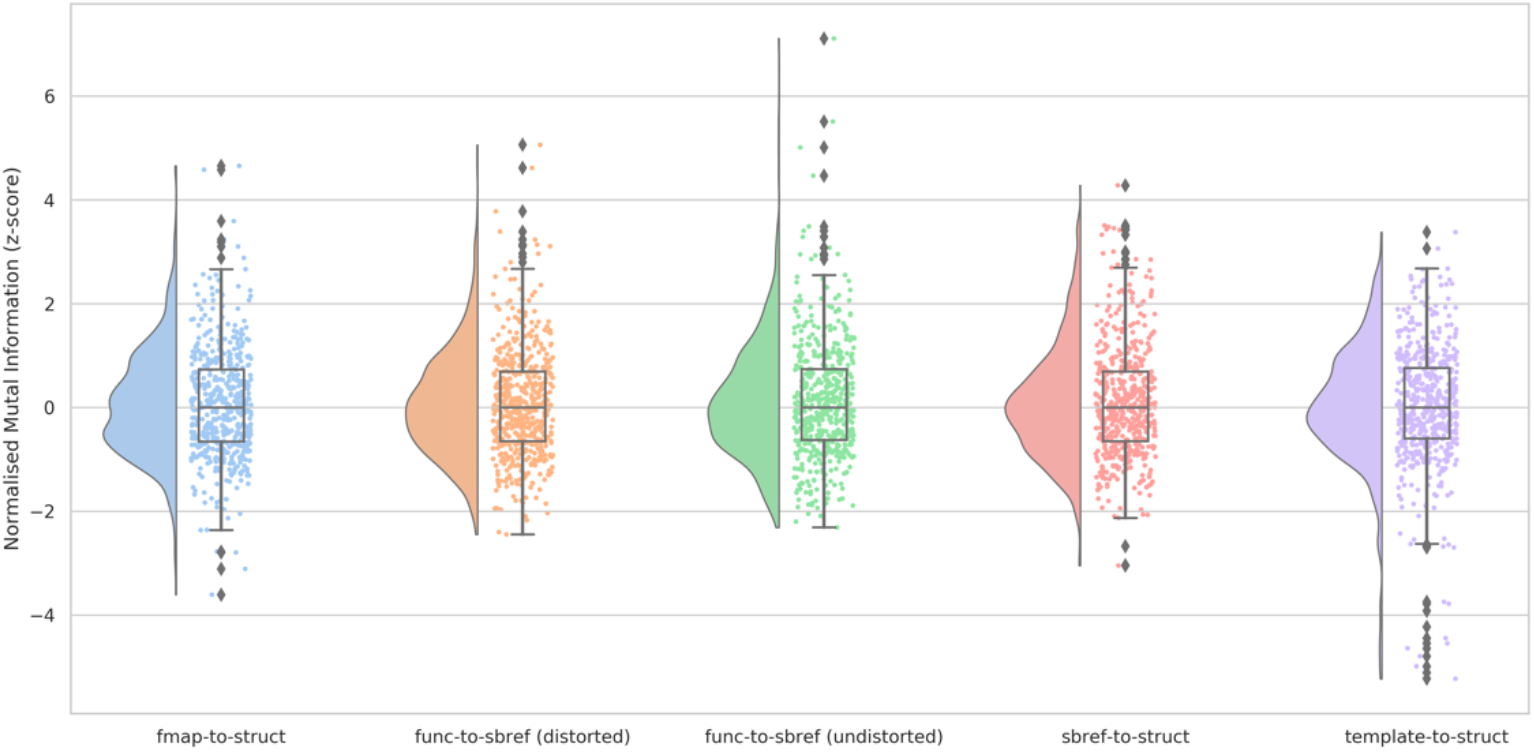
Distribution of the z-scored normalised mutual information between the source image and the reference image (both in reference space) for each of the primary registration stages fieldmap-to-structural, functional-to-sbref (distorted), functional-to-sbref (undistorted), sbref-to-structural, and template-to-structural. More positive NMI z-scores indicate more similarity and more negative NMI z-scores indicate less similarity.

Figure 4 presents example representative alignments of the fieldmap-to-structural, the sbref-to-structural and the standard-to-structural registrations, at differing levels of quality as quantified by the NMI similarity metric (see Supplementary Figure 9 for preterm examples). We selected the 5th, 50th and 95th percentile of NMI distribution, with the 5th representing the lower-end of alignment quality (whilst excluding outliers). The 5^th^ percentile fieldmap in this figure is a dual-echo-time-derived fieldmap magnitude and the lack of tissue contrast is clear; this not only makes registration to the structural space difficult, but also makes it hard to judge the quality of the registration. At the 50^th^ and 95^th^ percentiles, the fieldmap magnitude images are spin-echo-EPI-derived and have good tissue contrast and alignment to the structural space. The GM/WM boundary of the sbref-to-structural qualitatively appears to align well at all three percentiles; however, at the 5^th^ percentile there are clear anterior distortions which would impact the quantitative assessment of registration quality. Given that the sbref has been distortion corrected, we conclude that these distortions are irrecoverable signal loss. There are some observable alignment errors in the template-to-structural, most noticeably at the 5^th^ percentile. However, this appears qualitatively comparable to aligning adult data to a common template.

**Figure 4.**
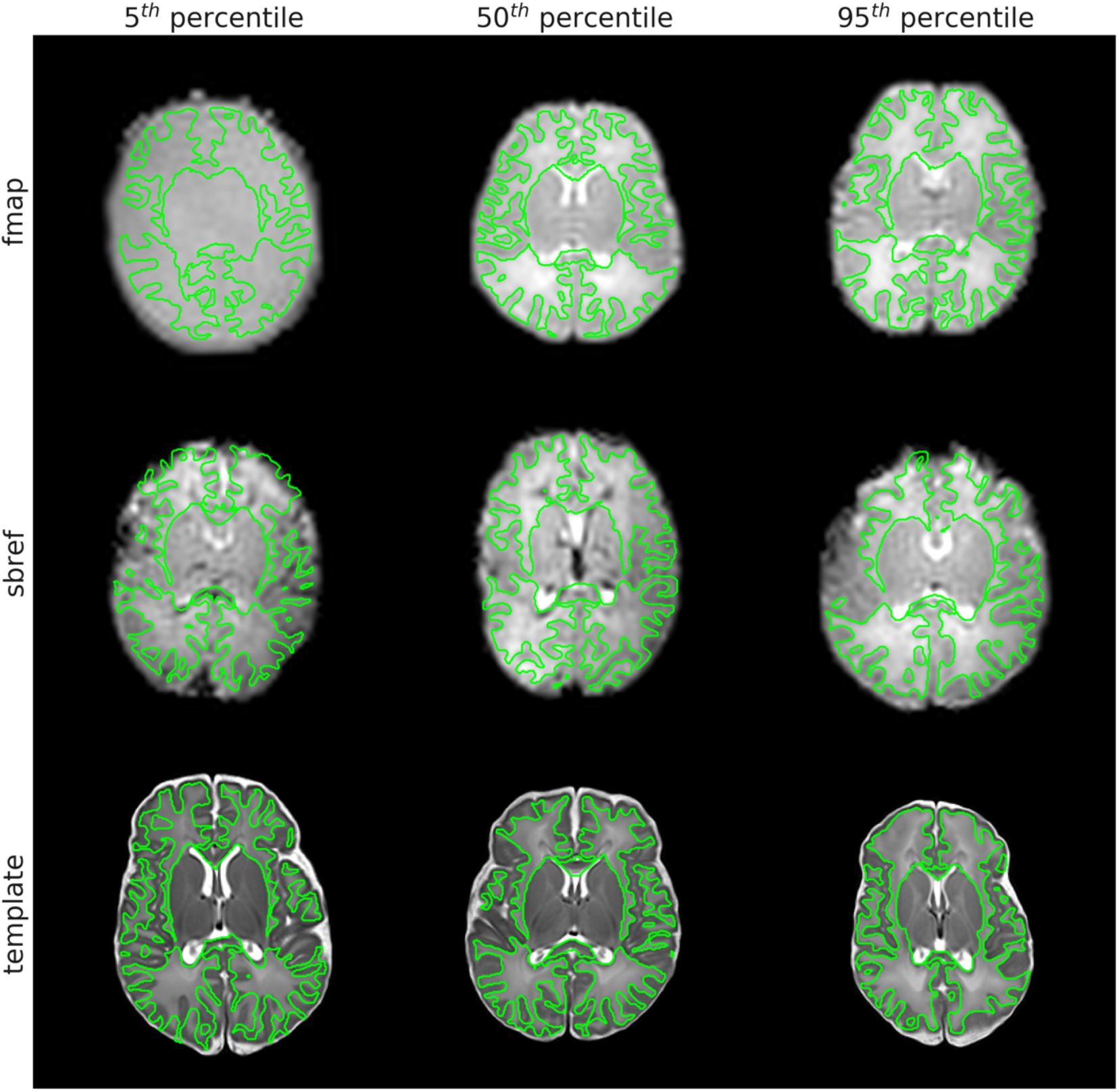
Examples of fieldmap, sbref, and template images resampled to the native structural reference space. The outline of the native structural white matter discrete segmentation is overlaid in green. Examples were selected at the 5th, 50th and 95th percentile of normalised mutual information between the source image and the reference image (both in reference space). Note: the 5^th^ percentile fieldmap is dual-echo-time-derived and therefore lacking tissue contrast, whilst the 50^th^ and 95^th^ percentile fieldmaps are spin-echo-EPI-derived

Figure 5 presents the dHCP 40-week T1w/T2w template, as well as group average and standard deviation across subjects of (1) the structural T2w in template space, and (2) the temporal-mean of the functional in template space. The group mean structural (T2w) has good anatomical contrast, although it is not as sharp as the template image, which likely reflects our decision to balance alignment with regularisation as discussed in Supplementary Section 9.3. The group mean functional also demonstrates anatomical contrast, but there are two distinct areas of lower intensity (observed in the mean) and high variability (observed in the stdev). The first is in inferior temporal and frontal areas which are most affected by susceptibility distortions and signal loss, and the second is in inferior occipital and superior-anterior cerebellum. It appears that this latter effect may be related to higher susceptibility induced variation (compared to adults) close to the transverse sinus in neonates, which is large in diameter and “ballooned” in the neonatal period as the venous system is still developing (Okudera et al., 1994). This observation is under further investigation.

**Figure 5.**
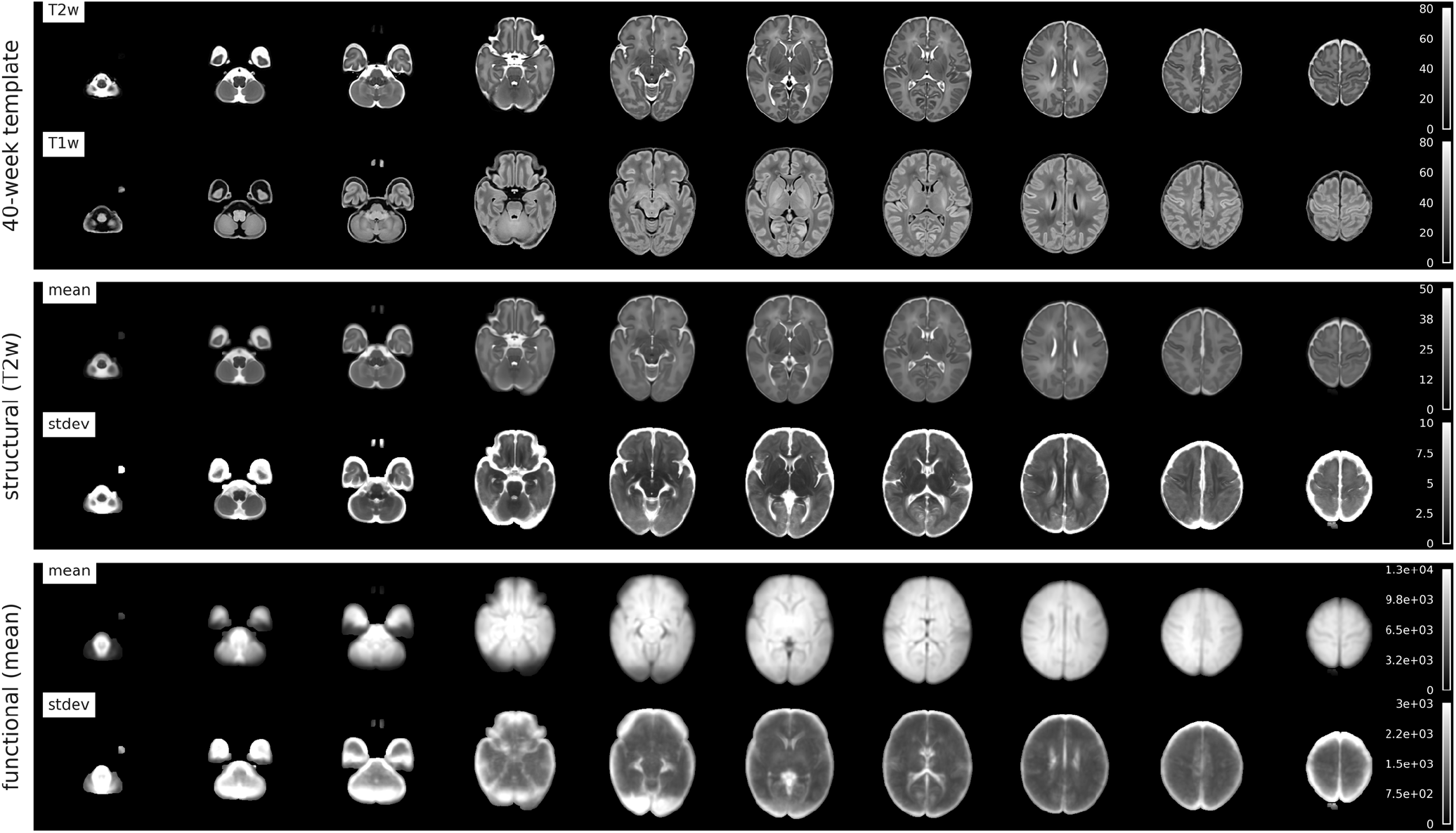
Upper: 40-week T2w and T1w dHCP templates. Middle: group mean and standard deviation (N=512) of structural T2w in template space. Lower: group mean and standard deviation (N=512) of functional (mean) in template space.

### 3.4. Susceptibility and motion correction

It is well documented that rfMRI analyses are very sensitive to subject head motion (Power et al., 2014). Head motion results in a number of imaging artefacts, many of which are not typically corrected in traditional adult pipelines, specifically:

1. **Volume and slice misalignment.** Volume misalignment is due to inter-volume movement and is usually effectively corrected using rigid-body registration-based motion correction. Intra-volume movement artefacts are a consequence of rapid subject movement during the sequential acquisition of the slices, or multi-band groups of slices, that constitute a volume. If, for example, the subject moves between the acquisition of the first and the second group of slices, the slices will no longer constitute a true volumetric representation of the object when stacked together, most noticeably by jagged edges of the brain (see Figure 6: Raw).
2. **Susceptibility-by-movement distortion.** Placing a subject in the scanner disrupts the static magnetic field because different tissues have different susceptibility to magnetisation. This field inhomogeneity results in distortions in the acquired image. The exact details of the disruption are defined by the configuration of tissue and air (sinuses, ear canals etc). To correct these distortions, it is common to estimate the field and use this to correct (unwarp) the acquired image. However, any subject movement that involves a rotation of the head around an axis non-parallel to the magnetic flux (z-axis) changes that field, which in turn changes the distortions in the image. That means that volumes acquired with the subject in different orientations will be subject to different distortions, and correction with a static estimate of the field, even with a rigid-body (re-)alignment, will not be sufficient to correct the changing distortions due to motion (see Figure 7: Rigid).
3. **Spin-history artefacts.** Movement during scanning can cause subsequent excitations to be misaligned with previous ones, resulting in differential excitation of magnetization at the slice boundaries; this leads to a striping effect in the image intensity (see Figure 9: Multi-band Artefact). The dHCP neonatal fMRI pipeline employs an ICA-based denoising method to remove spin history effects (see Section 3.5).

**Figure 6.**
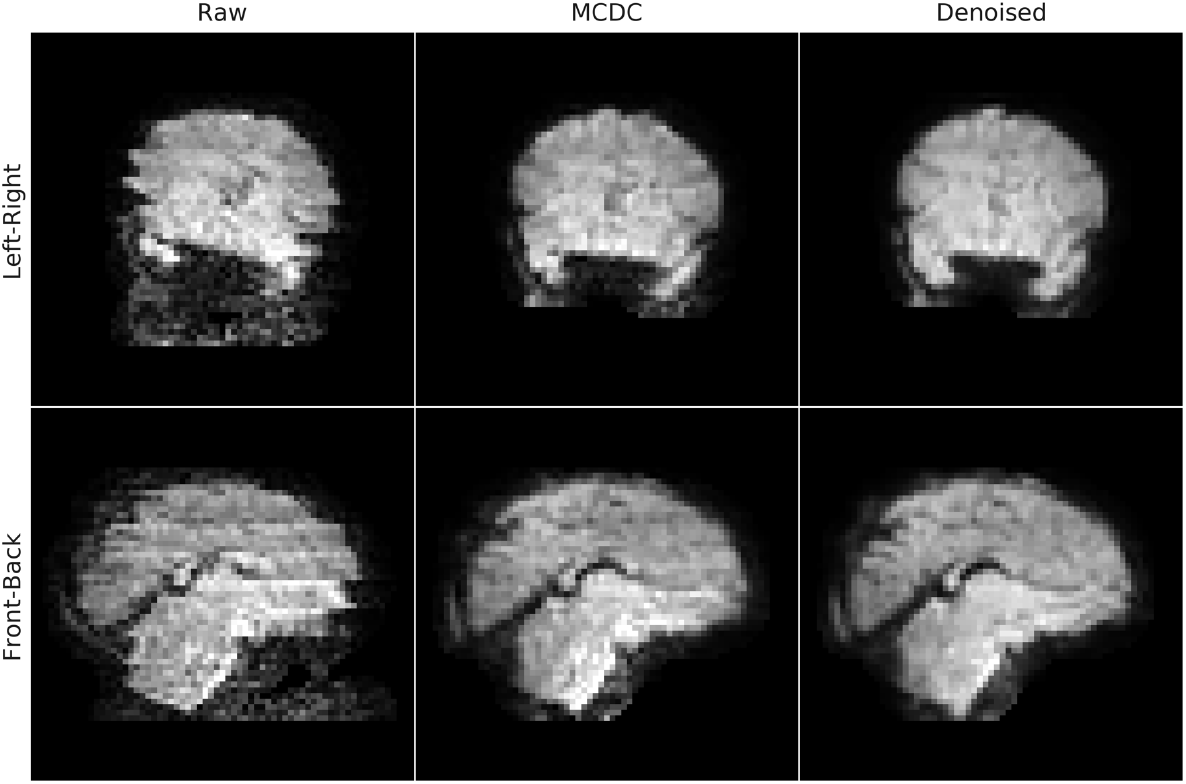
Exemplar single-volume of an EPI from a single-subject with intra-volume movement contamination from a leftright head movement (upper) and a front-back head movement (lower), before (Raw) and after motion and susceptibility distortion correction (MCDC), and after FIX denoising (Denoised).

**Figure 7.**
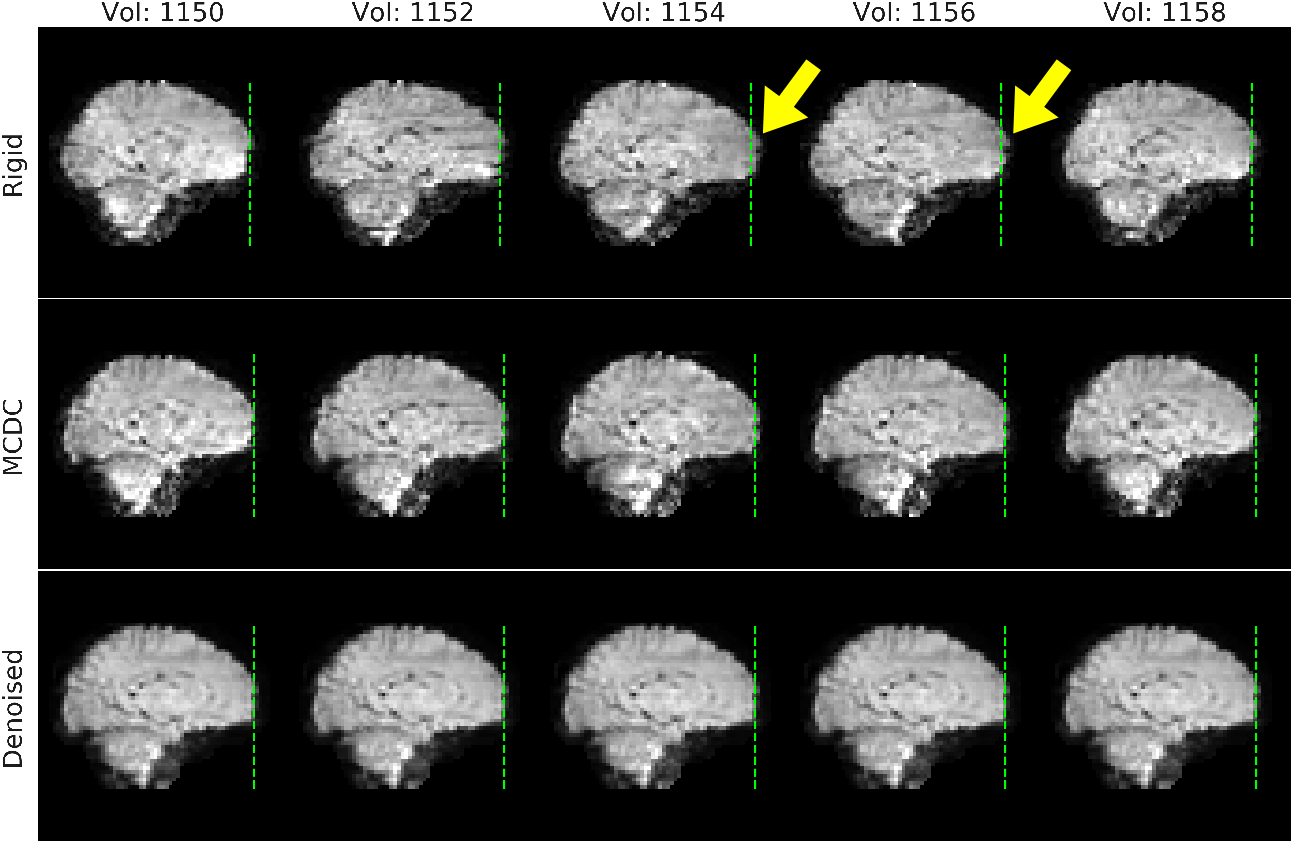
Five exemplar volumes of an EPI from a single-subject with susceptibility-by-movement distortion due to head motion. The rigid data in the top row have been rigid-body motion corrected, and anterior distortions can be observed in volumes 1154 and 1156 where the front of the brain extends beyond the reference line (green-dashed line). The anterior distortions are diminished after motion and susceptibility distortion correction (MCDC), and more so after denoising.

Spike regression (Satterthwaite et al., 2012) and scrubbing (Power et al., 2014, 2012), collectively referred as frame censoring, are popular and potentially effective alternative methods of dealing with head motion. We present a brief comparison of our approach with a frame censoring strategy in Supplementary Section 9.4. We have opted for a principled approach of correcting for artefacts introduced by motion (described in this section) combined with ICA-based denoising (see Section 3.5) which enables us to mitigate the effects of motion without excluding any subjects or time-points. Censoring methods remain a downstream option for researchers using the released data.

Motion and distortion correction (MCDC) are performed using the FSL EDDY tool. EDDY was designed for diffusion data and its extension for functional data is novel. When applied to fMRI, EDDY does not model eddy currents (as these are extremely low in fMRI), it instead treats each fMRI volume as a diffusion B0, using the temporal mean as a predictive model. The motivation for using EDDY on fMRI data is that it is capable of correcting for intra-volume movement artefacts (Andersson et al., 2017) and for artefacts associated with susceptibility-induced off-resonance field changes (susceptibility-by-movement artefacts) (Andersson et al., 2001).

EDDY performs a slice-to-volume (S2V) reconstruction to correct for intra-volume movement. This is achieved by using a continuous discrete cosine transform model of movement over time with degrees of freedom less than or equal to the number of slices (or multiband groups) (Andersson et al., 2017). EDDY corrects for the susceptibility-by-movement distortion (MBS) by estimating rate-of-change of off-resonance fields with respect to subject orientation (Andersson et al., 2018, 2001). These form parts of a Taylor-expansion of the susceptibility-induced field as a continuous function of subject orientation and allows for the estimation of a unique susceptibility field for each volume. The fieldmap in native functional space (fieldmap-to-functional, see Section 3.2) is input to EDDY and is used as the zeroth term of the Taylor-expansion of the field. The full MCDC proceeds by first estimating volume-to-volume movement, followed by estimation of slice-to-volume (intra-volume) movement. Finally, the changing susceptibility field is estimated, interspersed with updating of the slice-to-volume movement estimates. Once all the parameters have been estimated a single resampling of the data is performed using a hybrid 2D+1D spline interpolation (Andersson et al., 2017).

MCDC was evaluated on the dHCP-40 fMRI. For comparison, a rigid-body (between-volume) motion correction was also applied to the fMRI data using FSL MCFLIRT (Jenkinson et al., 2002a). Temporal signal-to-noise (tSNR) spatial maps were calculated for each subject on the raw (RAW) fMRI time-series, after rigid-body motion correction (RIGID), after slice-to-volume reconstruction (S2V), and after S2V and susceptibility-by-movement distortion correction combined (S2V+MBS). The tSNR maps, for each MCDC condition per subject, were resampled to standard space and voxel-wise group differences between the MCDC conditions calculated. Statistical evaluation of each of the tSNR difference maps was performed with a voxel-wise one-sample t-test using FSL RANDOMISE with 5000 permutations (Winkler et al., 2014). Thresholded group activity maps were corrected for multiple comparisons with false discovery rate (FDR) correction and a threshold of 1.67% (calculated as 5% divided by the number (3) of tests).

Rigid-body (RIGID) motion correction significantly improves tSNR compared to the RAW data (see Figure 8) mostly at the cortex and edges of the brain. S2V correction significantly improves tSNR compared to RIGID across the whole brain, and S2V+MBS further improves tSNR in anterior and posterior areas where susceptibility distortions are expected. The correction of intra-volume motion artefacts can be visually observed in an exemplar volume from a single subject in Figure 6 (MCDC), and the correction of susceptibility-by-movement distortions can be observed in an example subject in Figure 7 (MCDC). On the larger cohort, the combined motion and distortion correction stage (comprising S2V and MBS) significantly improves tSNR across the whole brain compared to the raw data (Figure 12).

**Figure 8.**
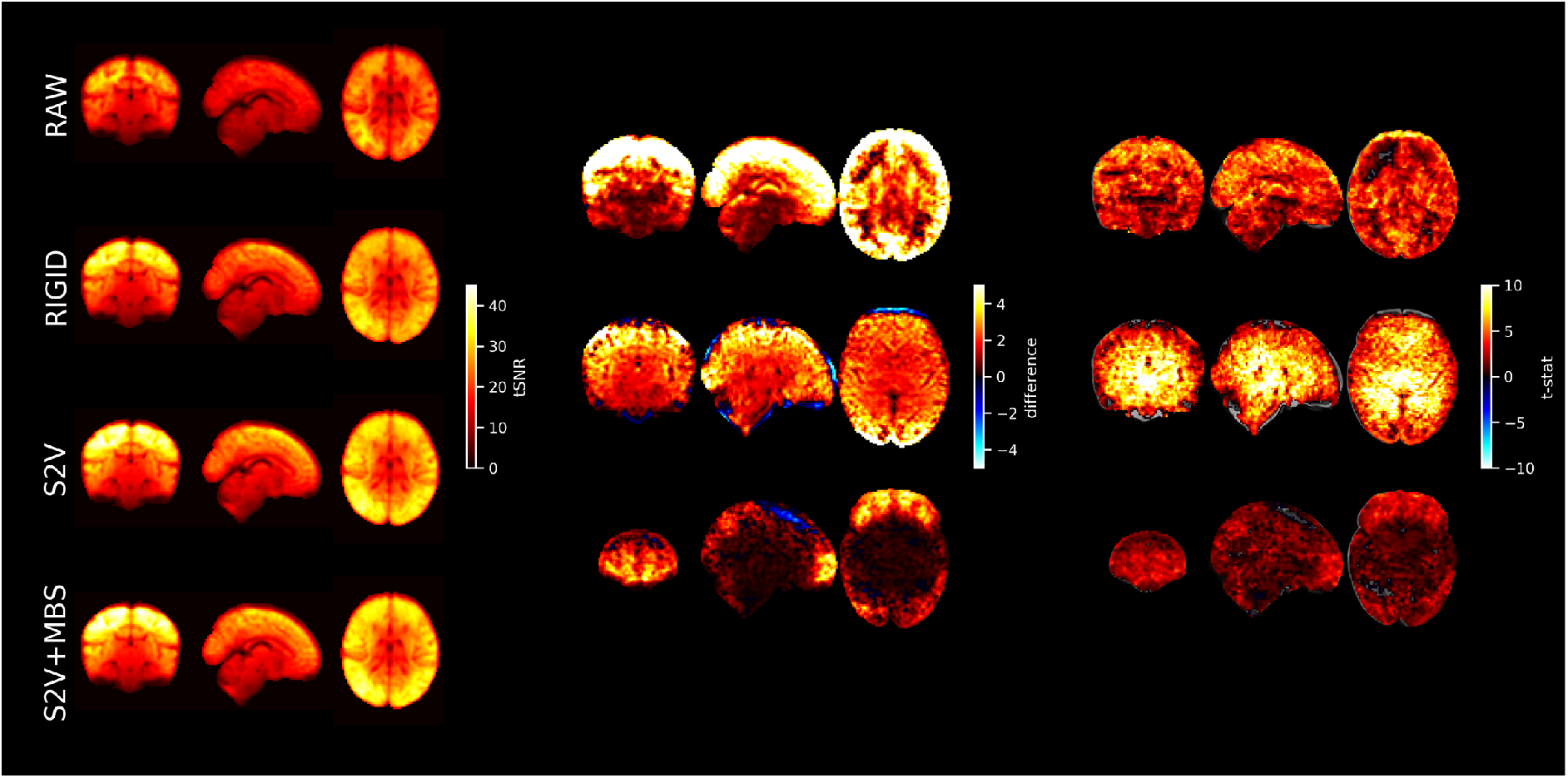
Left: mean tSNR (N=40) for raw EPI (RAW), rigid-body motion correction (RIGID), slice-to-volume motion correction (S2V), and S2V + susceptibility-by-movement distortion correction (S2V+MBS). Centre and right: difference maps and t-statistics for RIGID tSNR minus RAW tSNR (upper), S2V tSNR minus RIGID tSNR (middle) and S2V+MBS tSNR minus S2V tSNR (lower). Only significant results shown. Multiple comparison correction was achieved by FDR correction with a 1.67% threshold (5% divided by the number of tests). The slice coordinates for the difference maps and t-statistic maps were selected by the maximum t-statistic.

### 3.5. Denoising

Even after motion and susceptibility distortion correction there are still residual motion-related artefacts (for example, due to spin history effects) that need to be dealt with. There are also a number of additional structured noise artefacts, unrelated to head motion, that need to be addressed. Therefore, we perform a denoising procedure based on spatial independent component analysis (sICA) to remove these structured noise artefacts. SICA has proven to be a powerful tool for separating structured noise from neural signal and is widely used for denoising fMRI in both adults and infants (Alfaro-Almagro et al., 2018; Ludovica Griffanti et al., 2017; Mongerson et al., 2017; Smith et al., 2013) and has proven to be of great value in connectome projects including the (adult) Human Connectome Project and UK Biobank.

#### sICA

The (motion and distortion corrected) single-subject functional time-series was high-pass filtered (150s high-pass cutoff) to remove slow drifts, but no spatial smoothing was performed. SICA was performed using FSL MELODIC (Beckmann and Smith, 2004). The sICA dimensionality was automatically set using MELODIC’s Bayesian dimensionality estimation, however it was capped at an upper limit of 600 components (26% of the number of timepoints). Whilst we were conscious of not wanting to reduce the DOF too much, the decision to implement the cap was a pragmatic attempt to reduce the computational cost of the pipeline. We found in earlier iterations of the pipeline on smaller subsets of data that very few subjects were constrained by the 600 cap, and those that were constrained mostly contained a higher number of unclassified noise components (data not shown). It is also important to note that sICA is mathematically unable to separate global confounds (Glasser et al., 2019), however most of the confounds of interest (as detailed below) are spatially specific.

The main types of component observed in the dHCP sICA (see Figure 9 for examples) were:

1. **Signal**. Characterised by low frequencies and spatial clustering. Unlike adult ICs we often observe residual motion related jumps in the signal time-course, therefore we do not use that as a basis for exclusion.
2. **Multi-band artefact.** Characterised by the “venetian blind” effect (stripes in the sagittal and coronal planes) in the spatial maps and the time-course typically shows jumps that correlate with motion spikes. This artefact likely comprises the spin-history effects (described Section 3.4) as well as inter-slice leakage. Leakage results from imperfect multi-band reconstruction, and therefore residual signal from any given slice can “leak” to co-excited slices after separation, which results in correlations between the slices over time.
3. **Residual head-movement**. Characterised by a ring (or partial ring) at the edge of the brain in the spatial map, and time-course that strongly reflects the motion parameters or framewise displacement.
4. **Arteries**. In adults this would be characterised by activity in the spatial maps in the middle cerebral branches and a distinctive high-frequency spectrum. However, with the neonates we do not have sufficient spatial resolution and so rely almost exclusively on the power spectrum. This artefact is less commonly observed than the other artefacts in the dHCP data.
5. **Sagittal sinus.** Similar to adults, the main characteristic used to identify the sagittal sinus artefact is the superior inter-hemispheric ring in the sagittal plane of the spatial map. The sagittal sinus was often difficult to identify, and potential candidates were often labelled as “unknown” because the rater was not completely confident in the classification.
6. **Unclassified noise.** Does not clearly belong to one of the other structured noise categories and is characterised by a scattered spatial pattern, and often has jumps in the time course consistent with motion spikes.
7. **CSF pulsation.** Although not shown in the figure, we occasionally observe CSF pulsation characterised by a spatial overlap with the ventricles.

**Figure 9.**
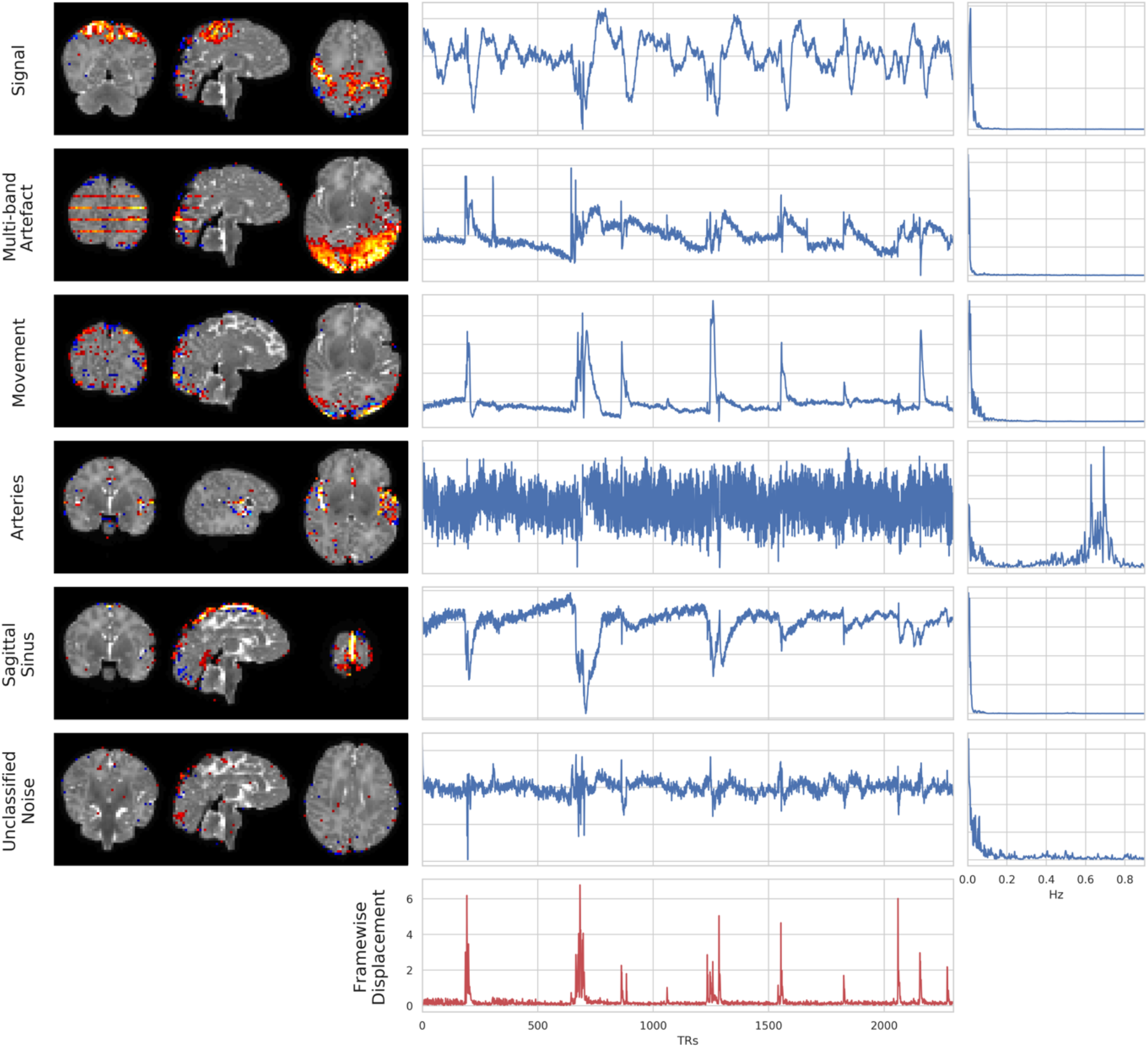
Exemplar spatial maps (left), time-courses (centre), and power spectra (right) for independent components (IC) from a single subject. Each row is a different IC that was manually classified as stereotypical for signal, multi-band artefact, head movement, arteries, sagittal sinus, and unclassified noise. Framewise displacement is plotted in the last row as a reference for the amount and timing of movement for this subject.

#### FIX

Artefactual independent components (ICs) were identified automatically using FMRIB’s ICA-based Xnoiseifier (FIX) v1.066 (Salimi-Khorshidi et al., 2014) which uses an ensemble machine learning classifier to label ICs as either **artefact** or **not** (ergo signal).

FIX was trained on a subset of manually labelled independent components (ICs) from 35 subjects. This subset was labelled using the scheme outlined in Griffanti et al. (2017) by a single investigator as signal, artefact, or unknown. The age range of the 35 subjects was 27.2 – 45.1 weeks PMA, the total number of manually labelled ICs was 4947. The manual labels for a subset of ICs from 5 subjects were double checked by two additional investigators with considerable experience of identifying artefacts in fMRI data (adult and neonatal), and who were following the specific guidelines given in Griffanti et al. (2017). The concordance of the primary rater with the first additional rater was 98.6%, and with the second additional rater was 100%.

As is the general theme of this paper, the nature of the neonatal data posed specific challenges for the manual IC labelling. In particular, the low spatial resolution relative to the size of the brain and the resulting partial-voluming made it more challenging to identify some of the components that would typically rely heavily on spatial features in adults, such as overlap with GM, WM and CSF. Therefore, we tended to rely more heavily on the timecourses. However, even in the time-courses, Griffanti et al. (2017) would recommend that signal should be “without sudden, abrupt changes” which we found to be too constraining in these data. Clearly stereotyped artefact components (head-movement, multi-band artefact, sagittal sinus, arteries, CSF pulsation, and unclassified noise) were labelled as “noise”. Any components that were unclear and/or difficult to judge were labelled as “signal”.

FIX was trained on the manually-labelled sICA data with a leave-one-out (LOO) training scheme. The median LOO true positive rate (TPR) was 100% and the median true negative rate (TNR) was 96%, indicating that the classifier erred on the side of inclusion (i.e. more likely to include noise than discard signal). This is a desired characteristic and is the reason that uncertain components were labelled as “signal”.

MELODIC/FIX was applied to the dHCP-538 dataset. The minimum number of ICs decomposed for a subject (i.e., ICA dimensionality) was 42, whilst the cap on the automatic dimensionality estimation of MELODIC resulted in 40 (7.4%) decompositions being constrained to the maximum dimensionality of 600. The proportion of ICs per subject/decomposition classified as noise and flagged for removal ranged from 53.2% to 100%, with a mean of 92.1% (see Figure 10). This is consistent with adults where the mean percentage of ICs classified as noise is typically ~70-90% (Ludovica Griffanti et al., 2017). The number of ICs per subject/decomposition classified as signal, and therefore flagged for retention, ranged from 0 to 46.8% with a mean of 7.9%.

**Figure 10.**
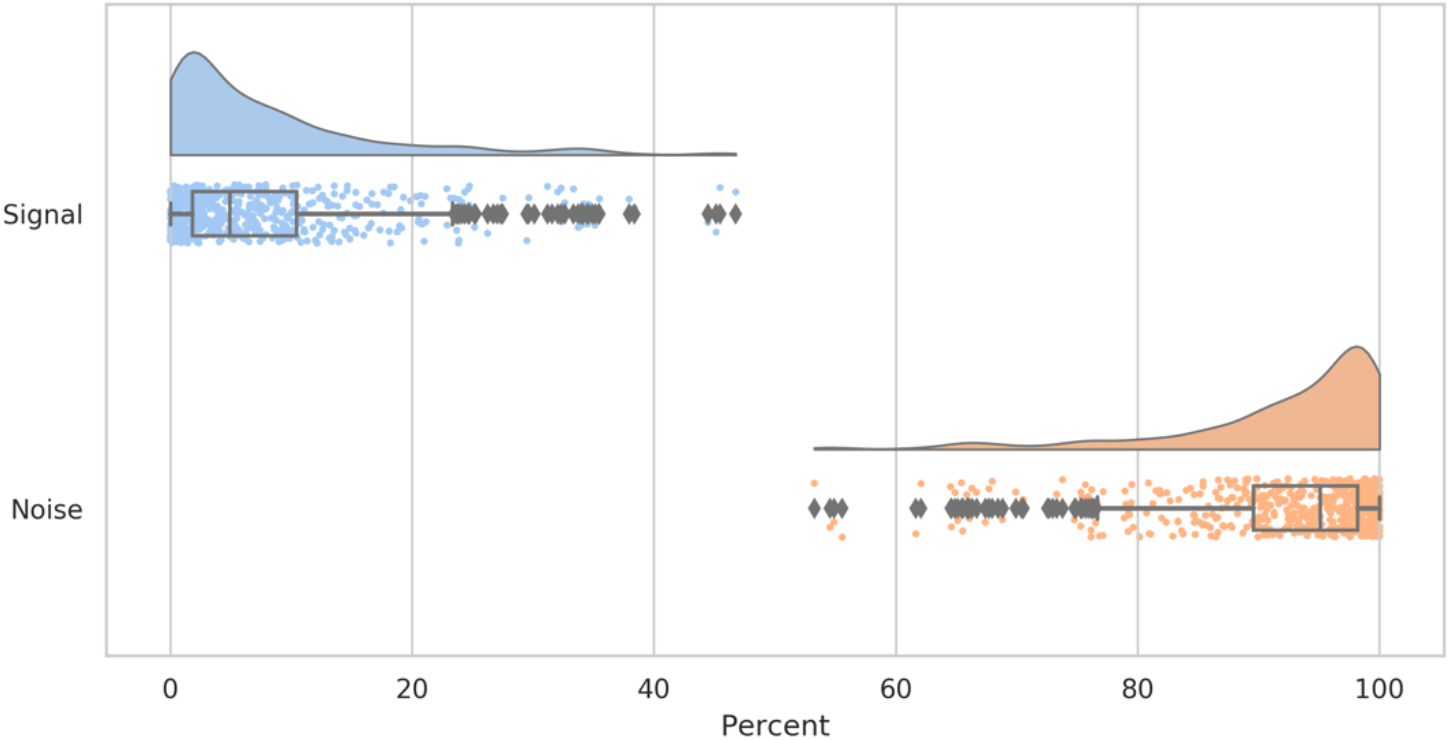
Distribution across decompositions of percentage of components (per decomposition) classified as signal or noise by FIX.

There were 19 subjects for whom all ICs were classified as noise. However, this does not necessarily mean that these data contained no signal, rather it means that signal information was not contained within the (constrained) reduced dimensionality on which the ICA was performed (and hence such signal would not be removed by the ICA denoising). Therefore, these data were still retained for further analysis.

Figure 11 shows that motion has a strong impact on the IC classification with the number of signal ICs decreasing with head movement (Spearman r=-0.5) and the number of noise ICs increasing (r=0.64). Age has a smaller impact with older babies tending to have more signal ICs (r=0.44) and less noise ICs (=-0.28). However, given that older babies tend to move more (see Supplementary Figure 7), and movement correlates with decreased signal classification, there could be an interaction where the true impact of age on signal classification is masked by the movement-signal effect, and vice-versa. Alternatively, the age-signal correlation may simply be a consequence of relative spatial resolution, with younger babies having smaller brains relative to the resolution of the acquisition

**Figure 11.**
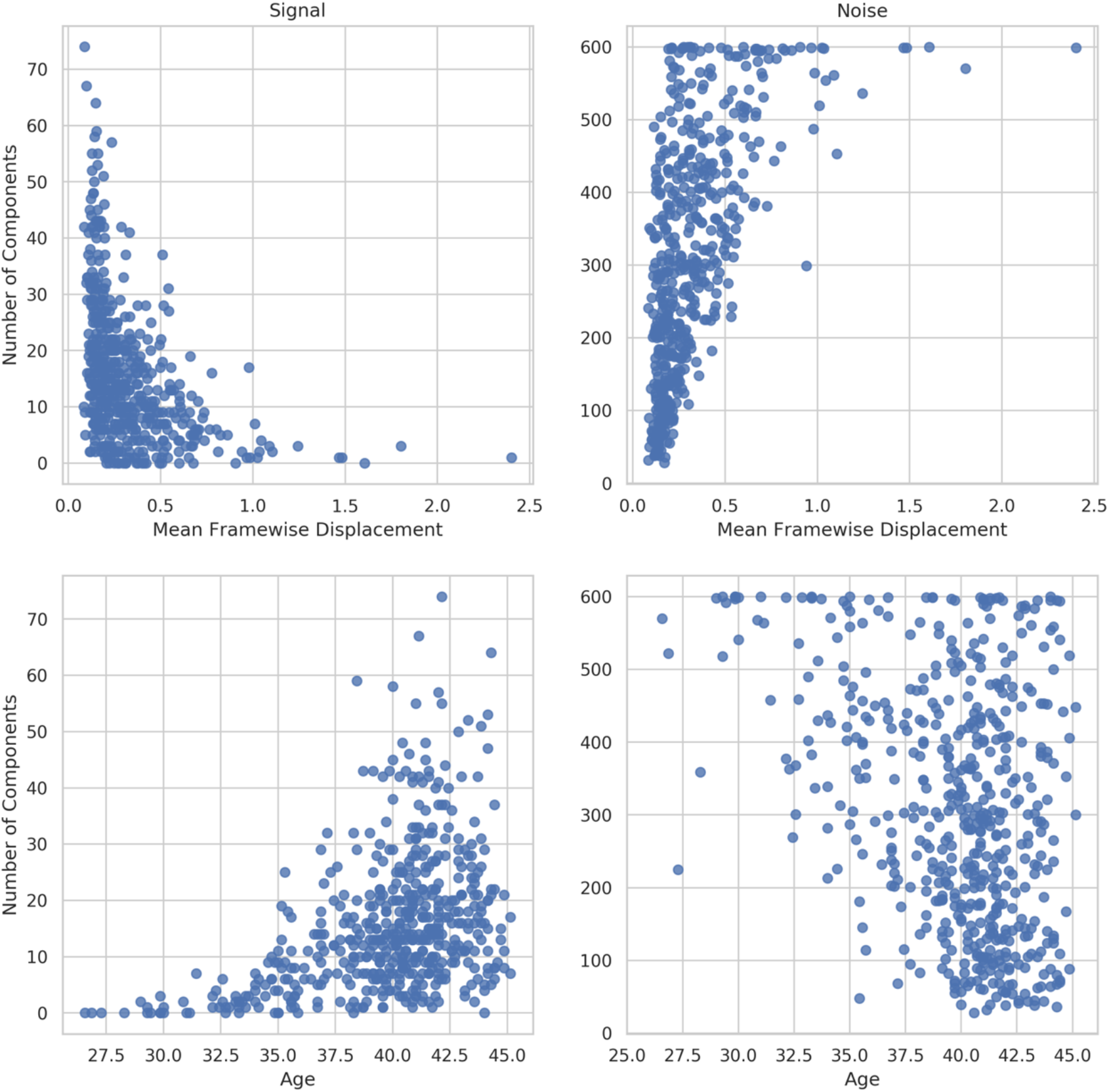
Correlation of the number of ICs classified by FIX as signal (left) and noise (right) with head movement, where mean framewise displacement is used as a surrogate for motion contamination. Age is the post-menstrual age-at-scan in weeks.

#### Nuisance regression

The FIX noise ICs and the motion parameters (see Table 3) were simultaneously regressed from the motion and distortion corrected functional time-series. The pipeline also supports inclusion of other nuisance regressors, such as FD outliers and DVARS outliers (see Table 3), physiological noise regressors, and tissue regressors. The inclusion of FD/DVARS outliers would effectively perform spike regression (Satterthwaite et al., 2013). However, given our objective to perform a *minimal* pre-processing we have not included additional nuisance regressors (beyond artefact ICs and motion parameters).

TSNR spatial maps were calculated for each subject on the raw (Raw) fMRI time-series, after motion and distortion correction combined (MCDC), and after denoising (Denoised) (see Supplementary Figure 9 for preterm examples). The tSNR maps were resampled to standard space and voxel-wise group differences between conditions calculated. Statistical evaluation of each of the tSNR difference maps was performed with a voxel-wise one-sample t-test using FSL RANDOMISE with 5000 permutations (Winkler et al., 2014). Thresholded group activity maps were corrected for multiple comparisons with FDR and a 2.5% threshold (calculated as 5% divided by the number of tests). After FIX denoising, the tSNR is substantially and significantly improved across the whole brain, with the greatest improvement seen in cortical areas (see Figure 12).

**Table 3.** Quality control metrics used in the dHCP fMRI pipeline.

| Metric | Description |
| --- | --- |
| Motion parameters (MP) | 6 rigid-body motion time-series (3 rotation, 3 translation) as estimated during motion correction |
| DVARS | DVARS is the RMS intensity difference between successive frames (Power et al., 2012) |
| DVARS outliers | Binarised DVARS with threshold = 75<sup>th</sup> percentile + (1.5 x inter-quartile range) |
| Framewise displacement (FD) | FD is calculated as the average of the rotation and translation motion parameter differences (Power et al., 2012) |
| FD outliers | Binarised FD with threshold = 0.25 mm |
| Temporal signal-to-noise ratio (tSNR) | Per-voxel temporal mean divided by the temporal standard deviation |
| Contrast-to-noise ratio (CNR) | Temporal standard deviation of the contrast divided by the standard deviation of the noise, where the contrast is the functional image minus the noise, and the noise is the residual of dual regressing the group spatial maps onto the functional image. |
| Normalised mutual information (NMI) | The normalised mutual information between a source image and a reference image (both in reference space). |

**Figure 12.**
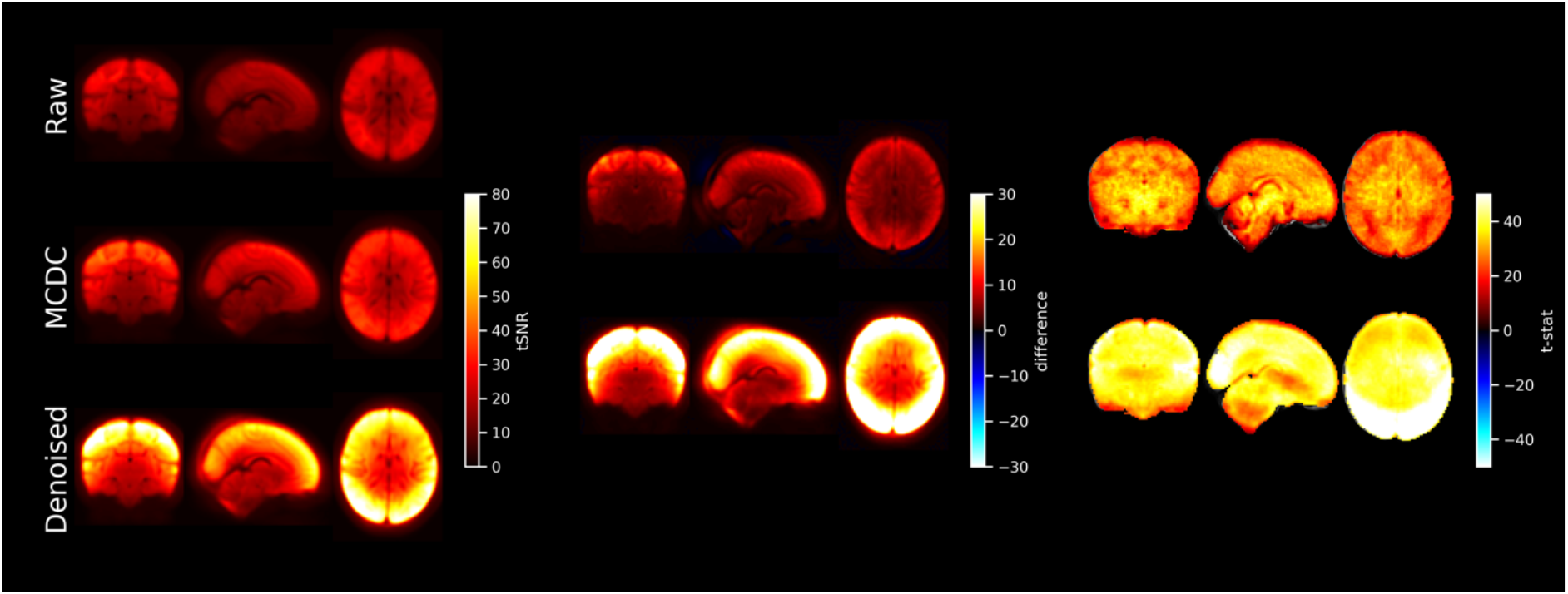
Left: mean tSNR (N=512) for raw EPI (RAW), motion and distortion corrected EPI (MCDC), and denoised EPI. Centre and right: difference maps and t-statistics for MCDC minus RAW tSNR (upper), and denoised minus MCDC tSNR (lower). Only significant results shown. Multiple comparison correction was achieved by FDR correction with a 2.5% threshold (5% divided by the number of tests)

TSNR informs us that there is less variance after denoising, but it does not delineate between noise variation or signal variation. Therefore, we additionally evaluate performance using spatial and network similarity to an unbiased group RSN template. Spatial and network matrix similarity to an unbiased group template was calculated for the raw (Raw) fMRI, after motion and distortion correction combined (MCDC), and after ICA+FIX denoising (Denoised). This involved running a group ICA (dimension=50) across all pre-processed data (all subjects; MCDC and Denoised) to generate group spatial maps that would not be biased to favour either MCDC or Denoised data in subsequent similarity calculations. The unbiased group maps were visually inspected and 13 RSN consistent maps identified (see Figure 13). All the unbiased group maps were then dual-regressed (Nickerson et al., 2017) onto the individual subject fMRI (Raw, MCDC, and Denoised) to yield subject-specific time-courses and spatial maps. Spatial similarity to the unbiased group template was then calculated as the spatial correlation between each subject-specific spatial map and the 13 unbiased group RSN maps. A network matrix was constructed for each subject by calculating partial correlation between all pairs of dual-regressed subject-specific time-courses. This resulted in a 13×13 matrix of partial correlations between all pairs of the 13 unbiased RSN-consistent group maps. The subject-specific network matrices were z-transformed and averaged (all subjects; MCDC and Denoised) to generate an unbiased group average network matrix. Network matrix similarity to the unbiased group average was then calculated as the correlation between each subject-specific network matrix and the unbiased group network matrix. The unbiased group RSN maps and group network matrix represent typical group output measures, and therefore effective pre-processing should increase similarity to the group templates.

**Figure 13.**
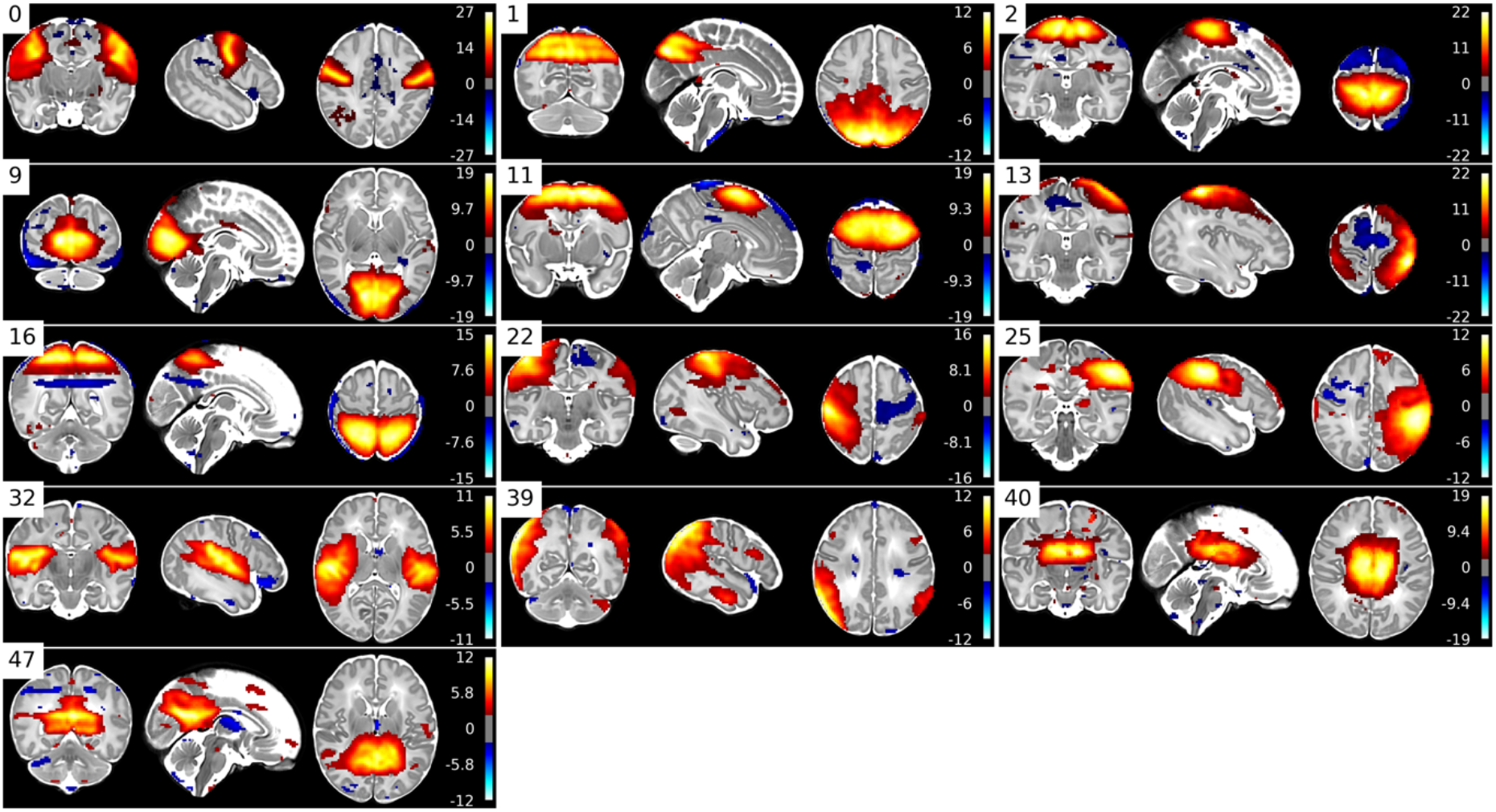
Unbiased group RSN template maps created from MCDC and Denoised data.

Group paired-differences in spatial and network similarity between (a) MCDC and raw, and (b) denoised and MCDC, were calculated using FSL RANDOMISE (Winkler et al., 2014) with 5000 permutations. Multiple comparison correction was achieved by FDR correction. Compared to Raw data, MCDC pre-processed data has significantly (p<0.025) greater spatial similarity to the unbiased group RSN maps in all of the 13 maps (see Figure 14), and significantly (p<0.025) greater network matrix similarity to the unbiased group network matrix (see Figure 15). Similarly, compared to MCDC pre-processed data, Denoised data has significantly (p<0.025) greater spatial similarity to the unbiased group RSN maps in 11 of 13 maps (see Figure 14), and significantly (p<0.025) greater network matrix similarity to the unbiased group network matrix (see Figure 15).

**Figure 14.**
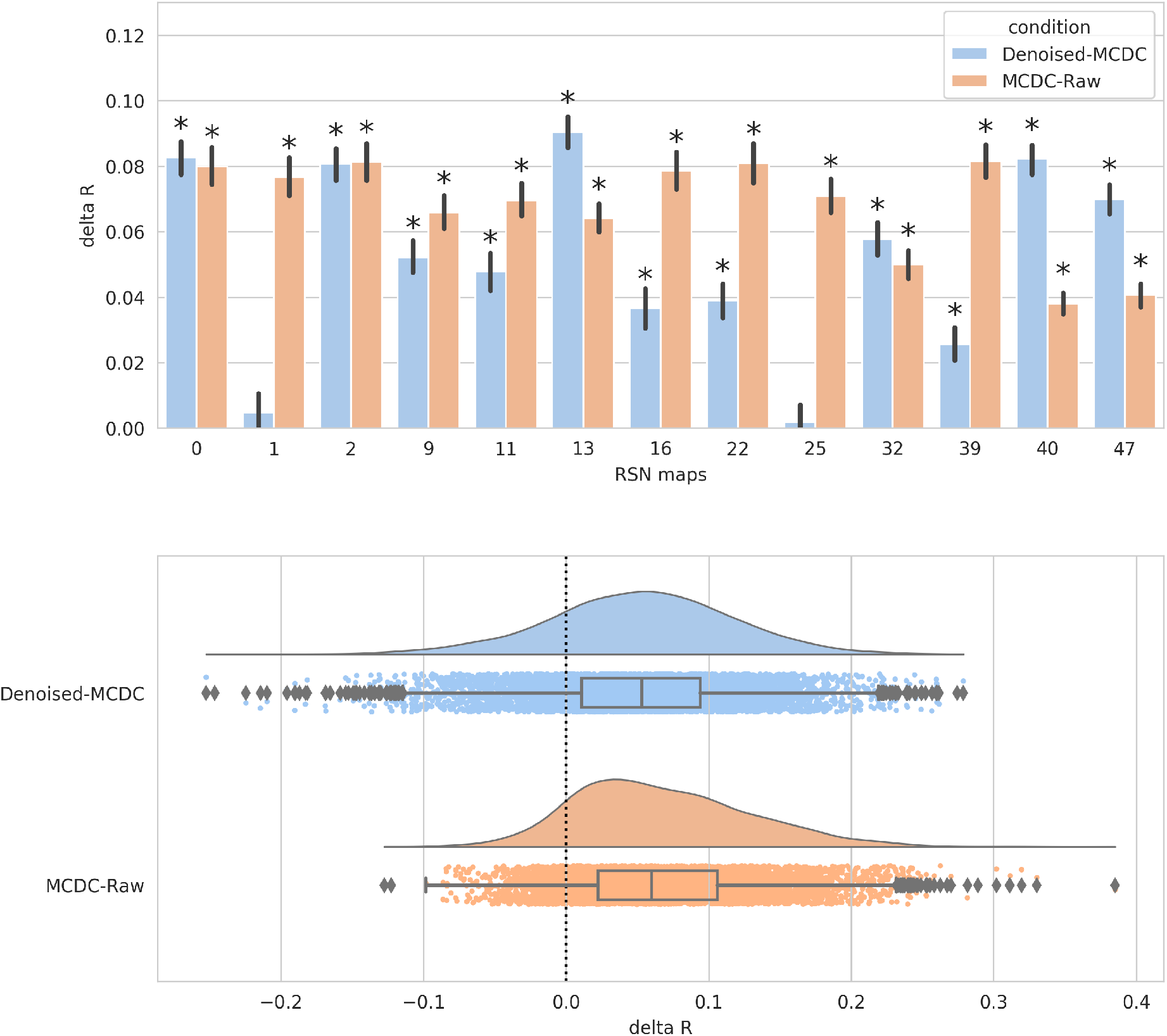
Mean paired-difference (Denoised-MCDC, and MCDC-Raw) of spatial similarity to the unbiased group template per map. Asterisks indicate significant differences. (Lower) Distribution of paired differences (Denoised-MCDC, and MCDC-Raw) of spatial similarity to the unbiased group template pooled over all spatial maps.

**Figure 15.**
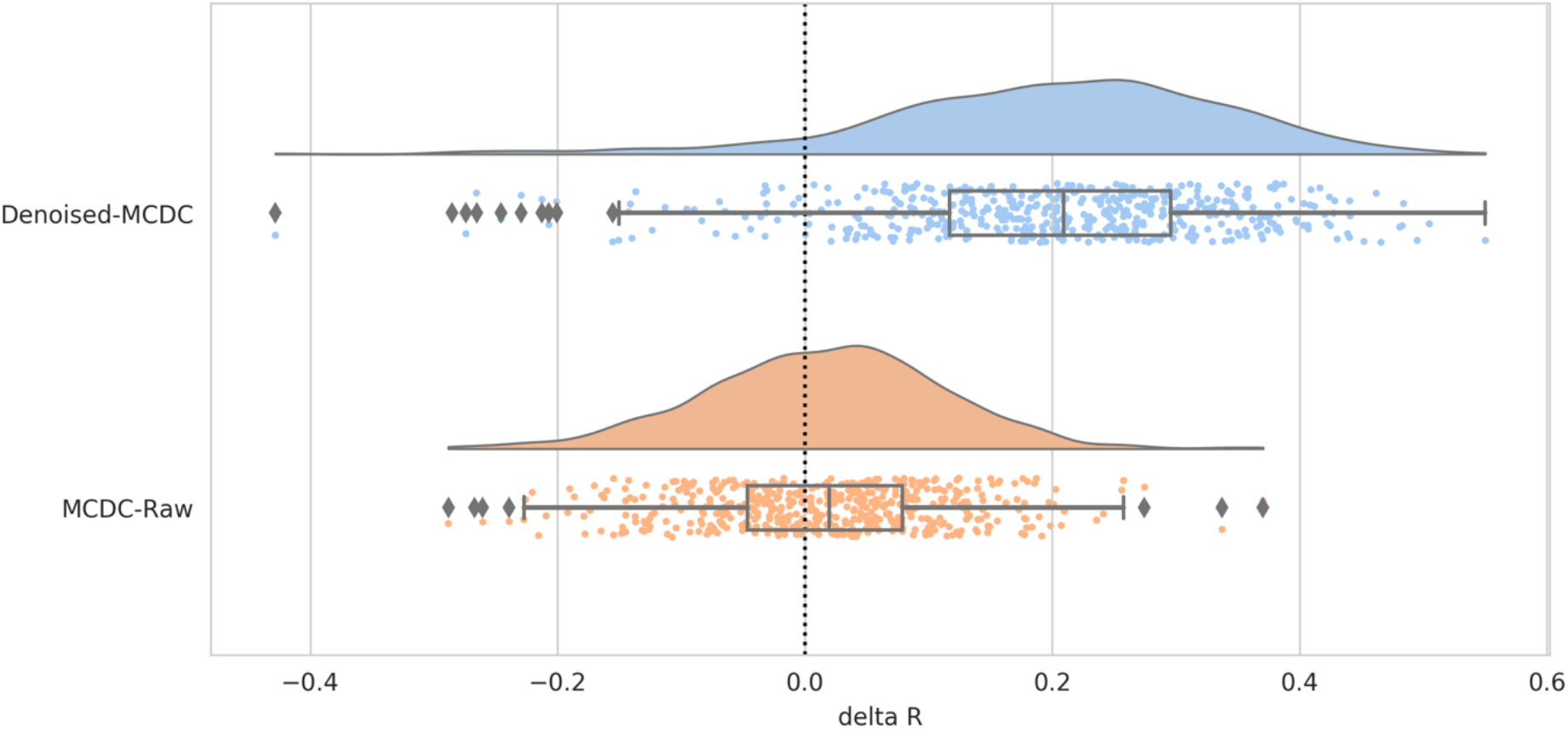
Distribution of paired differences (Denoised-MCDC, and MCDC-Raw) of network matrix similarity to the unbiased group network matrix.

These results suggest that the combination of motion and distortion correction with ICA+FIX-based denoising can substantially improve SNR by removing noise variance from the data, whilst at the same time maintaining signal and ultimately improving outcome measures.

Voxplots (aka. “carpetplots” and “grayplots”) comprise a heat-map of voxel x time fMRI intensities (with mean and linear trend removed) along with plots of nuisance time-series such as DVARS and framewise displacement (surrogates for motion). Voxplots were developed by Power (2017) and are advocated as an informative way to visualise and asses scan quality. We have adapted them by converting each heat-map to a z-score and using a diverging colormap so that it accentuates divergence from the mean of zero. Voxplots for a single example subject for raw, motion and distortion corrected (MCDC), and denoised fMRI are presented in Figure 16.

**Figure 16.**
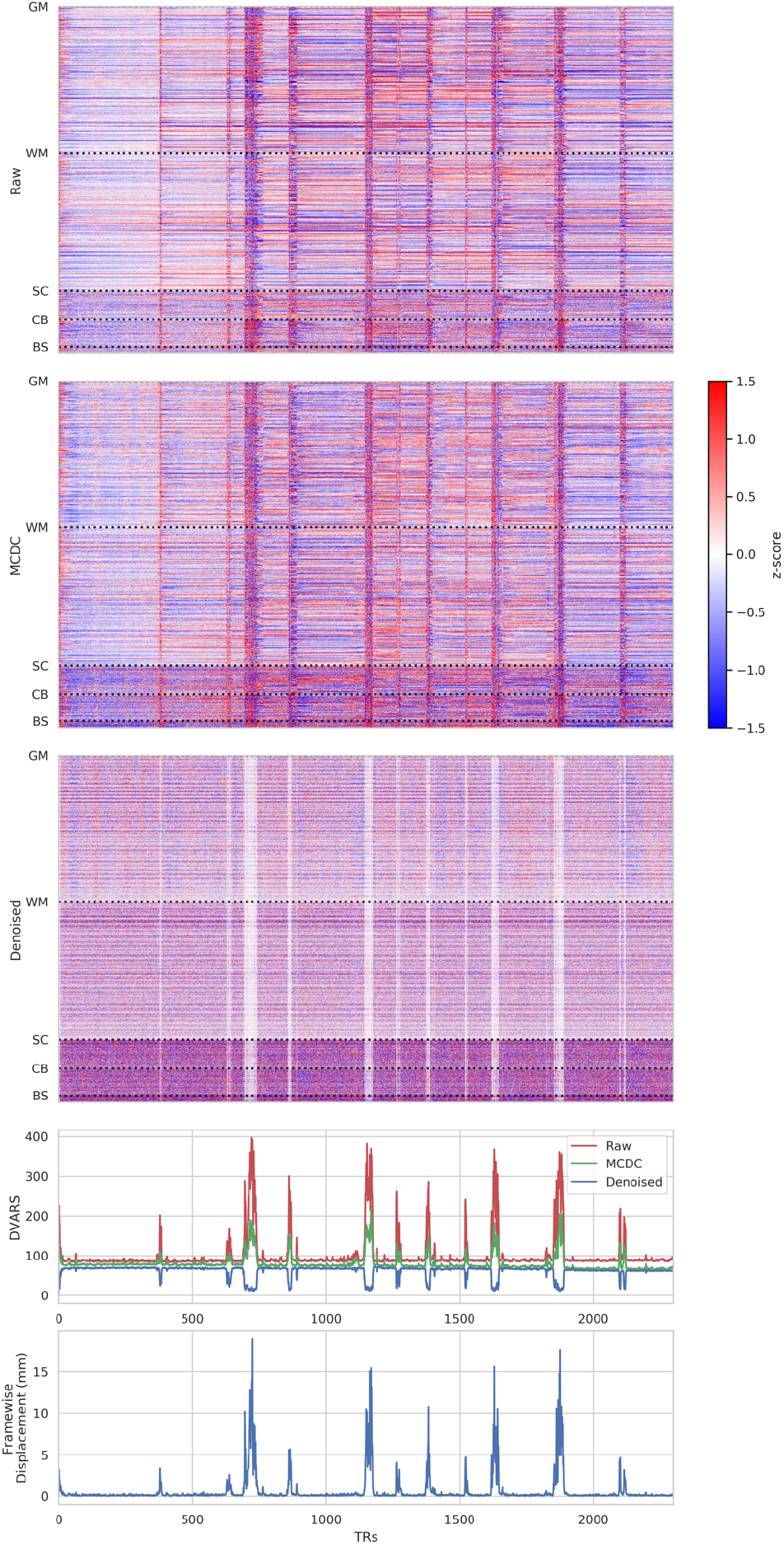
Voxplots for raw, motion and distortion corrected (MCDC), and denoised fMRI from a single exemplar subject. Voxplots are adapted from (Power, 2017) with the modification that each heat-map is converted to a z-score and a diverging colormap is used. Mean and trend were removed from each heat-map. GM=gray matter, WM=white matter, SC=sub-cortical, CB=cerebellum, BS=brainstem.

Strong vertical stripes can be observed in the pre-denoised data (raw and MCDC) that are contemporaneous with the worst spikes in the nuisance time-series (DVARS and framewise displacement).

After motion and distortion correction, there is less variation in-between the vertical stripes compared to the raw fMRI, which can also be observed in the DVARS plot as lower values between the major spikes. The heat-map also shows that this reduced variation is largely limited to the cerebral cortex and matter. The intensity of the vertical stripes also drops after motion and distortion correction, which is difficult to see in the heat-maps because each map is independently normalised to the standard score; however, it can be observed as lower amplitudes of the major spikes in the in the DVARS plot.

After denoising, there a several striking differences in the voxplot. Firstly, the strong vertical stripes are replaced with closer-to-zero vertical stripes, indicating that the denoising has removed much of the variation at these times and the remaining signal is closer to the mean. This is represented by dips in the DVARS plot during these times. If spike regression were used we would expect to see that these periods would be exactly the mean and would thus be zero in both the heat-map and DVARS plot. Thus, ICA-based denoising seems to provide a qualitatively similar result to spike regression in these periods of time when motion is the worst. Secondly, the tSNR difference between the white-gray-matter and the sub-cortical areas is further increased, consistent with the spatial group-maps presented in Figure 12.

ICA-based denoising can subsume the role of spike regression when the motion is severe, however, it is less aggressive and does not simply remove all information during these periods. In practise, it will often remove much of the variance, but it can in principle leave residual signal if it is not modelled as artefact. We consider that scrubbing and spike regression are a “hard” form of temporal censoring, whereas ICA-based denoising is a less aggressive “soft” spatio-temporal censoring. In either case, subsequent analyses must appropriately account for the noise removal approach and the reduction or removal of BOLD signal during high motion periods.

### 3.6. Quality control/assurance

The dHCP neonatal fMRI pipeline automatically calculates a number of QC metrics (see Table 3) and generates an HTML QC report for each subject (Supplementary Figure 10). The report presents the QC metrics for the individual within the context of group distributions for the corresponding metrics.

A subset of the metrics are converted to z-scores (using the median absolute deviation which is more robust to outliers than the standard deviation), and sign flipped as necessary so that more positive values are better and negative values are worse (see Figure 17). Subjects that score less than −2.5 on any of the subset of metrics are considered to have failed the QC and are flagged for further manual inspection. The specific subset of measures used are mean denoised DVARS, mean denoised tSNR, func-to-sbref NMI, sbref-to-structural NMI, structural-to-template NMI, and fieldmap-to-structural NMI. Under this regime, out of the total of 538 subjects, 5 subjects failed mean denoised DVARS, 4 failed denoised tSNR, 3 failed sbref-to-struct NMI, 18 failed template-to-struct NMI, and 4 failed fmap-to-struct. However, there was overlap of subjects between these failures, with a total of 26 subjects failing and thus 512 subjects passing. It is anticipated that a large proportion of the failed subjects can be recovered with improvements to the template-to-struct registration that are currently under investigation.

**Figure 17.**
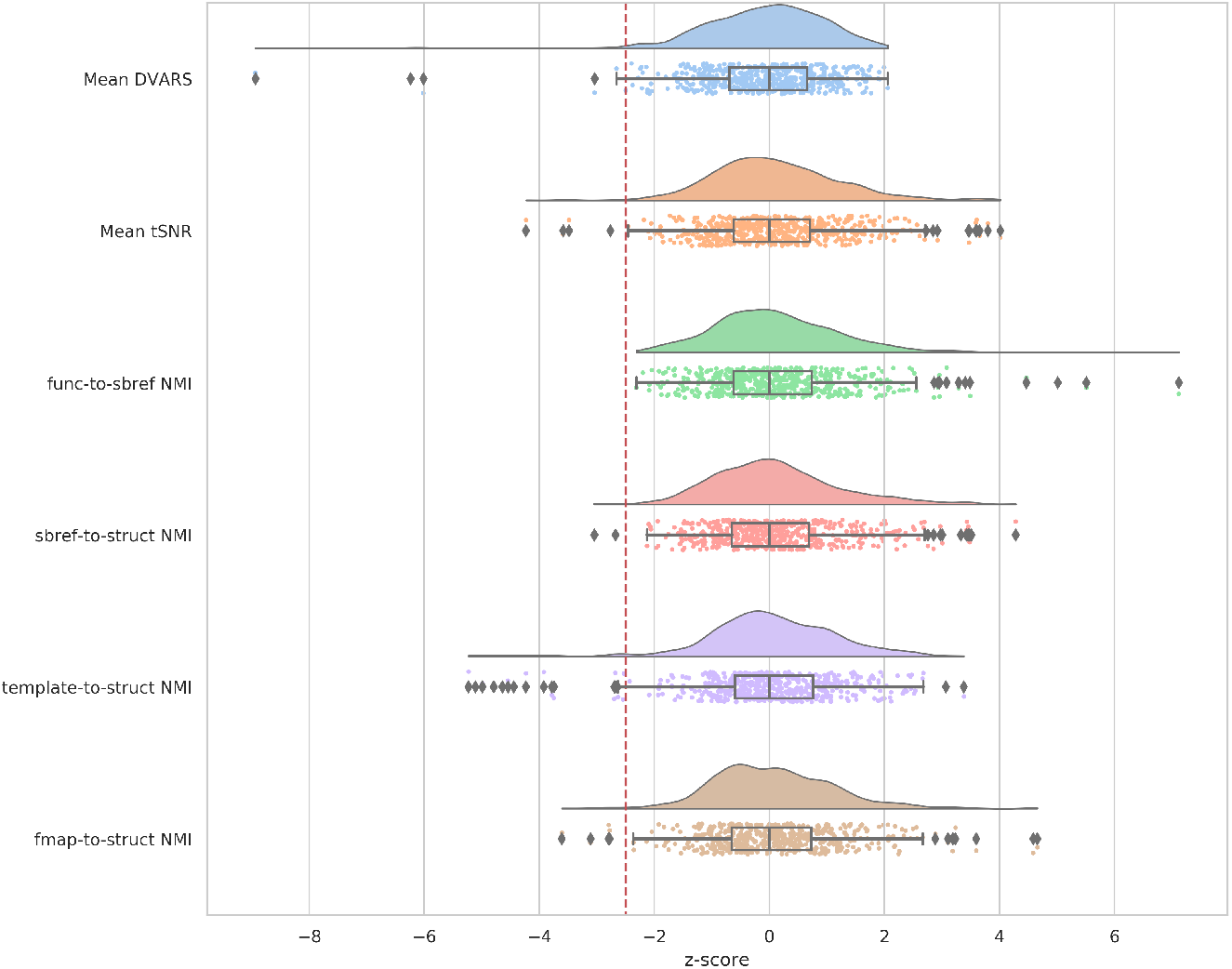
Group z-distributions of QC metrics. More negative z-scores indicate poorer quality on the respective metric. Z-scores less than −2.5 (indicated by red dashed line) are flagged as failing the pipeline and require further inspection.

### 3.7. Resting-state networks

Here we present group RSNs and their association with PMA as a validation that the dHCP neonatal fMRI pipeline can identify plausible RSNs, and to demonstrate the granularity of what can be extracted from this challenging cohort with suitable pre-processing.

For the derivation of resting-state networks, we use PROFUMO, an implementation of the probabilistic functional modes (PFM) model as defined in Harrison et al. (2015). This approach uses a hierarchical Bayesian model to decompose the data into a set of functional modes (i.e., RSNs). Unlike most ICA-based approaches, group and subject-specific spatial maps associated with these modes are estimated simultaneously. The PFM model includes a number of different terms that regularise the decomposition, including, for example, hierarchical priors that encourage consistency in both the spatial layout of RSNs across subjects, and their patterns of functional connectivity (i.e., connectomes). The model also takes haemodynamics into account, as these dominate the temporal characteristics of the BOLD signal. As such, we base these priors on the work of Arichi et al. (2012), who quantify haemodynamic responses for neonates. Specifically, we used the FSL FLOBS tool (Woolrich et al., 2004) to construct a term and pre-term HRF model using the reported haemodynamic response characteristics described in Arichi et al. (2012). Supplementary Figure 11 depicts the term and pre-term HRFs that we constructed, along with the default adult HRF used in PROFUMO. Only a single HRF model can be used within a PROFUMO analysis, therefore we used the term HRF model as the majority of the dHCP neonatal cohort are term age. The term HRF is characterised by a longer time to peak positive amplitude, a smaller positive peak amplitude and a deeper negative undershoot period relative to the adult HRF (T. Arichi et al., 2012).

Formally, the PFM model simultaneously decomposes the dataset from each subject *s* and run *r*, *D*^(*sr*)^ into a set of *M* modes. These consist of subject-specific spatial maps *P*^(*s*)^, along with run-specific amplitudes *h*^(*sr*)^ time courses *A*^(*sr*)^ and network matrices *α*^(*sr*)^. These can be combined into a matrix factorisation model at the subject level i.e. *D*^(*sr*)^ ≈ *P*^(*s*)^ × *diog*(*h*^(*sr*)^ × *A*^(*sr*)^. Note that for the data analysed here, all subjects only have one run of fMRI data; we therefore refer to all parameters as subject-specific from now on.

These subject-level decompositions are linked via a set of hierarchical priors, which represent the group-level description of the data (and include, amongst others, the group-mean spatial maps and connectomes). This forms a complete probabilistic model for the data, and the group- and subject-level information is inferred together via a variational Bayesian inversion scheme. This is covered in more detail in Harrison et al. (2019).

In summary, PROFUMO not only infers subject-specific information in a sensitive manner, but it also infers the group-level properties of the RSNs themselves. The analysis was performed on the denoised volumetric data of 512 dHCP subjects that passed the pipeline QC (see Section 3.6) to resolve resting-state networks (RSNs). Prior to PROFUMO the data were spatially smoothed (FWHM=3mm) using FSL SUSAN, with an intensity threshold of 75% of the contrast between the median brain intensity and the background (Smith and Brady, 1997), and normalised to the grand median intensity as per FSL FEAT (Jenkinson et al., 2012). 16 RSNs were identified (see Figure 18) that show good correspondence to both adult (Smith et al., 2013) and infant RSNs (Doria et al., 2010; Mongerson et al., 2017). We also regressed the RSNs on age and demonstrate significant changes in shape and amplitude of these networks with development from 29-45 weeks post-menstrual age (see Supplementary Section 9.5). See Supplementary Figure 9 for examples of dual-regressed single-subject spatial maps from preterm subjects.

**Figure 18.**
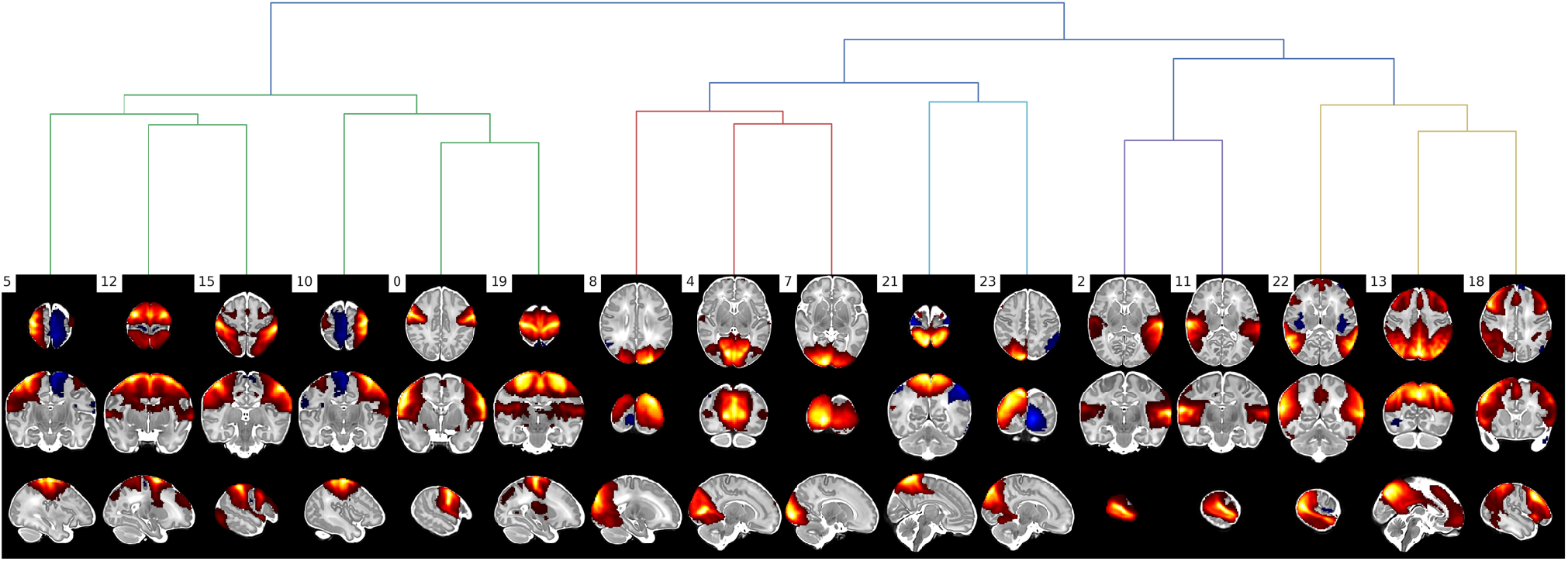
PROFUMO modes qualitatively assessed as corresponding to adult resting-state networks. Hierarchical clustering based on spatial correlation between the modes Warm colours are positive, and cool colours are negative.

## 4. Discussion

In this paper we present an automated and robust, open source pipeline to generate high-quality minimally pre-processed neonatal fMRI data. The pipeline was developed to preprocess the neonatal rfMRI data from the dHCP project for open-access release to the neuroimaging community. Using this pipeline on the dHCP neonatal cohort, we have been able to resolve 16 resting-state networks with fine spatial resolution that are consistent with adult networks from the Human Connectome Project. Furthermore, we have sufficient spatiotemporal granularity to demonstrate significant changes in shape and amplitude of these networks with development from 29-45 weeks post-menstrual age.

A strong motivation during development of the pipeline was that it should perform “minimal” pre-processing to ensure that the scientific community would not be restricted in the subsequent analysis that they can perform on the data (although the fully raw data are also being made available). To this end, we have sought to minimise pre-processing, particularly resampling, whilst maximising output quality. The pre-processing steps we have included were selected and evaluated to ensure they provided a robust and principled approach to mitigating the specific challenges of neonatal data. The pipeline was run on 538 subjects from the dHCP neonatal cohort and only 26 failed due to quality control restrictions. These failures were mostly due to poor registration of the structural T1w/T2w to the template/atlas space. This is discussed further below.

Automation was another key motivator in developing the pipeline. Given the number of subjects to be scanned as part of the dHCP project it was important to minimize manual intervention as much as possible. To this end, the pipeline is entirely automated including quality control and reporting. Operator intervention is required at only two stages, 1) to manually label a subset of independent components to train FIX, and 2) to visually inspect cases flagged by QC as outliers. Furthermore, the FIX classifier trained on dHCP data will be released along with the pipeline, which means that the first manual intervention step may be avoided if one’s data are sufficiently similar to the dHCP acquisition data. However, users of the pipeline should perform a thorough evaluation if applying the dHCP trained FIX classifier to other cohorts.

Subject head motion is the most challenging confound observed in the dHCP neonatal cohort. This motion disrupts the BOLD signal and can result in slice misalignments, susceptibility-by-movement distortions, and spin history artefacts. Such artefacts are not typically dealt with in existing fMRI pre-processing pipelines. We present a novel application of the EDDY tool, which was originally designed for diffusion data, to correct for the slice misalignments and susceptibility-by-movement distortions. We further incorporate an ICA-based denoising procedure to remove spin-history effects and any residual motion artefacts. This ICA-based denoising can also account for a variety of other artefacts including multi-band artefacts, arteries, CSF pulsation, and sagittal sinus. As a consequence of these pre-processing strategies, we see large and significant improvement in tSNR across the whole brain, but particularly in cortical areas. This improvement is driven largely by a reduction in variation from the aforementioned artefacts.

An important consideration that needs to be made when using the pipeline is that EDDY can only be run using a GPU. The EDDY-based motion and distortion correction on a single dHCP subject (2300 volumes) takes 6-12 hours on a NVIDIA K80 GPU. If limited resources mean that EDDY is not viable, the pipeline can fallback to a (CPU-based) rigid-body volume-to-volume registration-based motion correction and static fieldmap-based distortion correction. Substantial improvements in speed have already been achieved in EDDY, and we expect the version in the next release to be 3-4 times faster. Furthermore, an fMRI-specific version of EDDY is intended to be released in a future version of FSL.

The dHCP neonatal fMRI pipeline includes a robust registration framework to align the functional data with both the subject structural space (T2w) and the group standard space. Registration is made challenging by the rapidly changing size and gyrification of the neonatal brain, and the variable contrast caused by maturing myelination and the fast-multi-band EPI sequence. The protocol uses a series of primary registrations which can then be combined into composite transforms to move between the target spaces. Care was taken to achieve high-quality primary registrations so that alignment errors would not accumulate when creating the composite registrations. To this end, the BBR cost function was found to be superior for intra-subject registrations and was used wherever possible. The weakest link in the registration protocol was the template-to-structural non-linear registration, which resulted in 18 subjects being excluded by the automated QC due to insufficient alignment quality. Ongoing work is examining improvements to this registration step, including using the GM probability as another registration channel, as per Makropoulos et al. (2018), and optimising the parameters of the registration tools.

The dHCP neonatal fMRI pipeline is not intended to be a single static release just for processing the dHCP data. During development we have focussed on flexibility and generalisability of the pipeline beyond the dHCP data. We have evaluated the pipeline on non-dHCP neonatal task fMRI data (Baxter et al., 2019), and collaborators within our centre have been evaluating the pipeline on non-neonatal data. To coincide with this paper, the first version of the pipeline will be released publicly (https://git.fmrib.ox.ac.uk/seanf/dhcp-neonatal-fmri-pipeline) and this will mark the beginning of what we plan to be an ongoing, open, and hopefully collaborative, development process. To this end there is a roadmap of future features that are either already under development or planned:

1. We have developed bespoke methods (adapted from adult HCP pipelines) for mapping the fMRI data to the surface and writing out to CIFTI format. These methods are still under evaluation.
2. An fMRI-specific version of EDDY is under development that will incorporate a model that is better suited to fMRI data. Significant speed-ups have already been achieved and will be part of the next FSL release.
3. Improvements to template-to-structural registration.
4. Partial BIDs derivatives support is implemented, but full support is planned.
5. Extensions for task fMRI have been developed and tested and will be merged into the pipeline

The second dHCP data can be obtained from the dHCP website (http://www.developingconnectome.org/second-data-release). This website also contains detailed release notes (http://www.developingconnectome.org/release-notes) that describes each file and will allow users to cross-reference this manuscript to the data.

## 5. Conclusion

We have presented an automated and robust pipeline to minimally pre-process highly confounded neonatal data, robustly, with low failure rates and high quality-assurance.

Processing refinements integrated into the dHCP fMRI framework provide substantial reduction in movement related distortions, resulting in significant improvements in SNR, and detection of high quality RSNs from neonates that are consistent with previously reported infant RSNs (Doria et al., 2010; Mongerson et al., 2017). Ongoing analyses are probing the fine structure of these networks, and their variability across subjects and age, with the aim of defining a multi-modal time-varying map of the neonatal connectome. The scientific community will be able to apply this pipeline to explore their own neonatal data, or to use publicly released dHCP data (pre-processed with this pipeline) to explore neonatal connectomics.

## 7. Acknowledgements

We are grateful to Dr Mark Chiew for advice on multi-band artefacts, and Assoc. Prof. Kenneth Pope for assistance with the neonatal HRF. The research leading to these results has received funding from the European Research Council under the European Union Seventh Framework Programme (FP/20072013)/ERC Grant Agreement no. 319456. We are grateful to the families who generously supported this trial. The work was supported by the NIHR Biomedical Research Centres at Guys and St Thomas NHS Trust. The Wellcome Centre for Integrative Neuromaging is supported by core funding from the Wellcome Trust (203139/ Z/16/Z). TA is supported by a MRC Clinician Scientist Fellowship [MR/P008712/1]. JA is supported by the Wellcome Trust Strategic Award (098369/Z/12/Z) and by the NIH Human Connectome Project (1U01MH109589-01 and 1U01AG052564-01). MJ is supported by the National Institute for Health Research (NIHR) Oxford Biomedical Research Centre (BRC). JOM is supported by a Sir Henry Dale Fellowship jointly funded by the Wellcome Trust and the Royal Society (Grant Number 206675/Z/17/Z). SRF is supported by the NIH Human Connectome Project (1U01MH109589-01 and 1U01AG052564-01). LB was funded via a Medical Research Council Studentship. SJH was supported by the grant #2017-403 of the Strategic Focus Area “Personalized Health and Related Technologies (PHRT)” of the ETH Domain. JB is supported by the European Regional Development Fund under grant KK.01.1.1.01.0009 (DATACROSS).

## 9. Supplementary

### 9.1. Fieldmap comparison

Visual inspection identified that 12.7% (75 of 590) spin-echo EPI in the dHCP data was contaminated in all volumes, such that picking the best pair of spin-echo EPI volumes was inadequate. In this circumstance, the fall-back procedure was to use the dual-echo-time-derived fieldmap instead of the spin-echo-EPI-derived fieldmap. To ensure that the dualecho-time and spin-echo-EPI derived fieldmaps could be used interchangeably, we evaluated the similarity between the two. For each subject the spin-echo-EPI and dual-echo-time fieldmaps were resampled to the native functional space and then converted to a voxel displacement/shift map using FSL FUGUE. The shift maps were then masked by an eroded brain mask, to avoid edge effects as a consequence of registration misalignment. Supplementary Figure 3 presents an example dual-echo-time and spin-echo-EPI derived fieldmap from a single subject, as well as the distribution of voxel displacements for all inbrain voxels from 409 subjects that had good quality dual-echo-time and spin-echo-EPI derived fieldmaps. The single-subject fieldmaps look qualitatively similar, although the dualecho-time-derived fieldmap appears smoother and the spin-echo-EPI-derived fieldmap appears to have greater values. This is supported by the voxel displacement distribution where the spin-echo-EPI-derived fieldmap has a slightly greater mean voxel displacement than the dual-echo-time-derived fieldmap. Two factors likely contribute to this difference, 1) the dual-echo-time fieldmap was acquired at lower resolution than the spin-echo-EPI (3×3×6mm and 2.15mm isotropic respectively), and 2) the dual-echo-time-derived fieldmap was low-pass filtered as part of the reconstruction process. Furthermore, the distribution of the voxel-wise difference between the dual-echo-time and spin-echo-EPI displacement/shift maps shows that 95% of voxels differ by less than 1 voxel shift and 99% by less than two voxels. We also inspected the distribution of spatial correlation between dual-echo-time and spin-echo-EPI derived fieldmaps across subjects, and observed good correspondence with 75% (i.e., 25^th^ percentile) of subjects showing correlation > 0.6. Given that the ground truth is unknown and that both the dual-echo-time and spin-echo-EPI derived fieldmaps are qualitatively and quantitatively similar, we felt justified in using the dual-echo-time-derived fieldmap as a back-up in cases where the spin-echo-EPI-derived was excessively contaminated by movement.

**Supplementary Figure 1.**
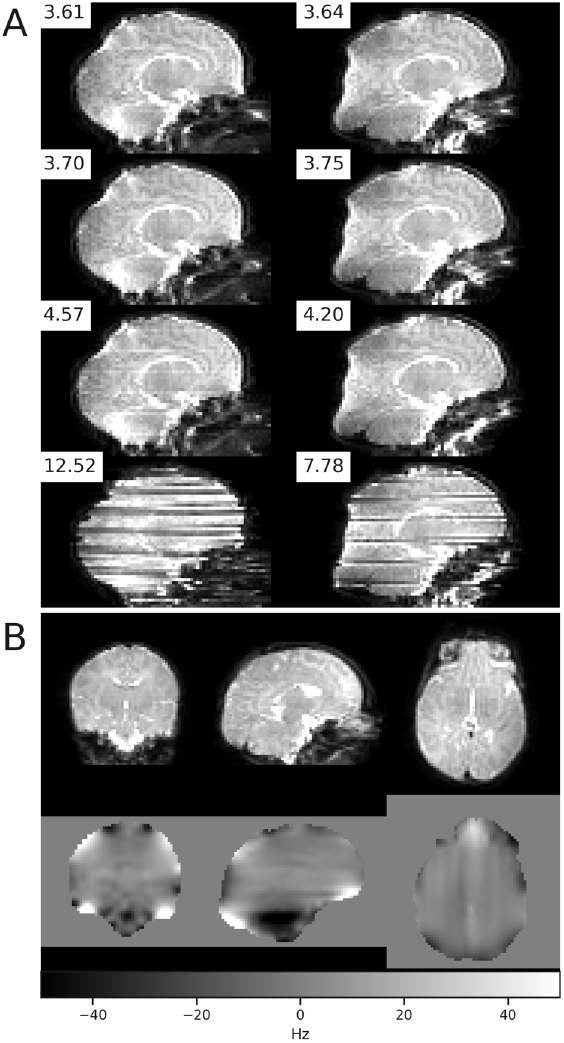
(A) Eight volumes of the spin-echo EPI from a single subject with AP (left) and PA (right) phase encode directions. Z-smoothness scores are presented with each volume. The two volumes in the last row have stereotyped striping artefact due to subject movement, resulting in higher z-smoothness scores. The two volumes in the first row were selected as the two “best” images based on z-smoothness. (B) Motion and distortion corrected spin-echo EPI (upper) and estimated susceptibility-induced off-resonance field (lower) derived from the spin-echo EPI in (A) using FSL TOPUP.

**Supplementary Figure 2.**
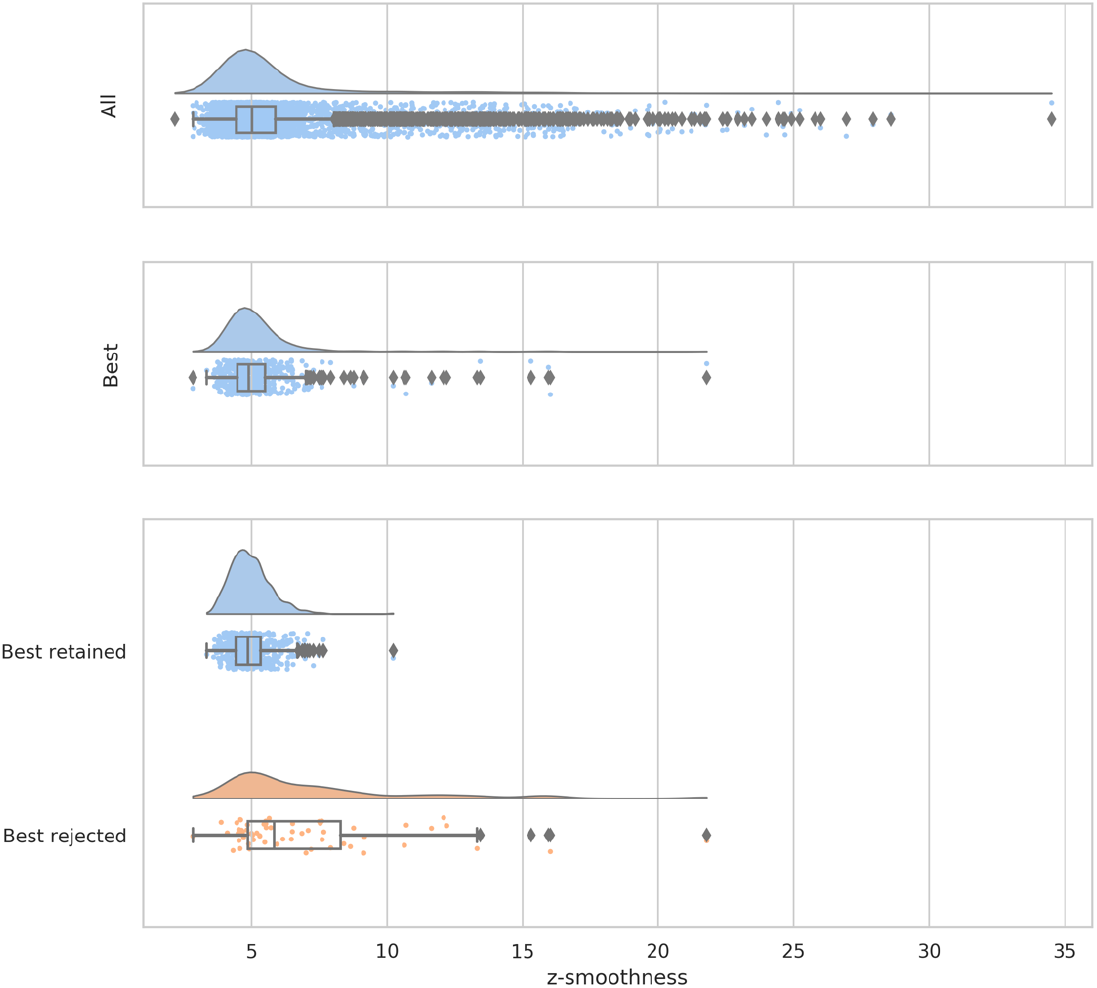
(Upper) Distribution of z-smoothness across all spin-echo-EPI volumes (8 per subject) across all subjects (N=538). (Middle). Distribution of z-smoothness for selected “best” quality spin-echo EPI volume across subjects. Best is defined by, first, picking the best pair of spin-echo-EPI volumes (one per phase encode direction) that had the lowest z-smoothness within subject, and then using the maximum z-smoothness value from that pair. (Lower) Same distribution as the middle plot but split by whether all spin-echo-EPI volumes were retained or rejected by visual inspection.

**Supplementary Figure 3.**
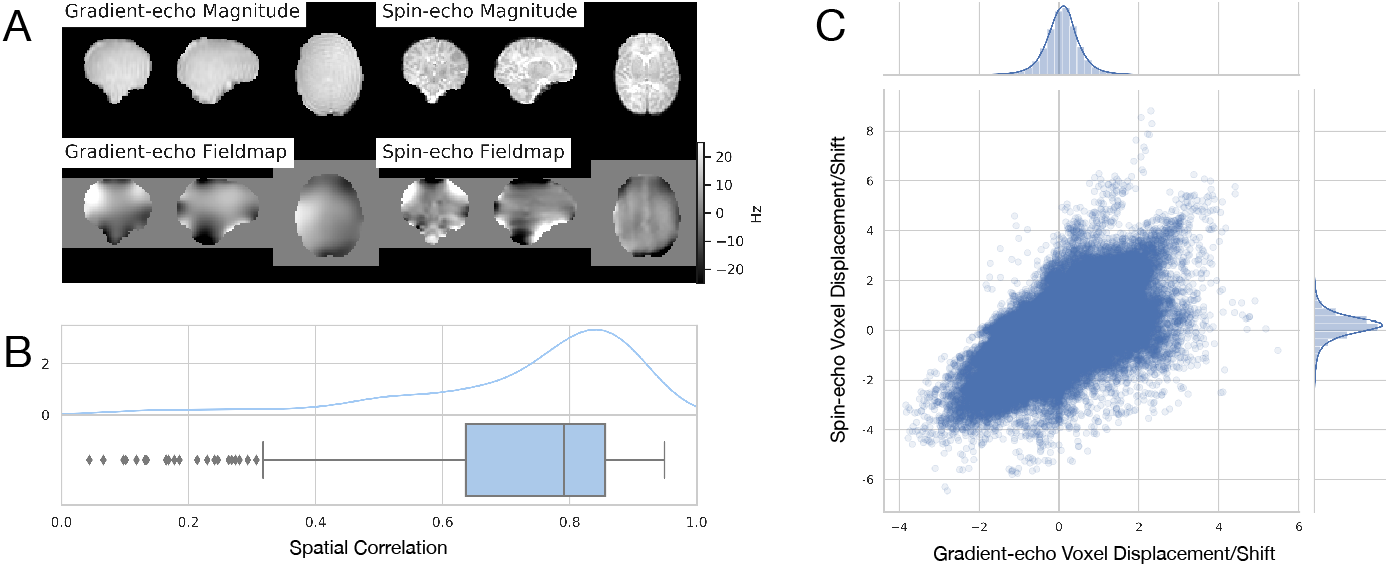
(A) Exemplar gradient-echo and spin-echo derived fieldmaps and magnitude from a single-subject. The spatial correlation of the two fieldmaps is 0.71. (B) Distribution of spatial correlation between gradient-echo and spin-echo fieldmaps from 409 subject. (C) Distribution of voxel displacement/shift for in-brain voxels from 409 subjects. To improve visualisation of the scatter plot density, only a 10% random sample of the (>11 million) voxel displacements/shifts are plotted.

### 9.2. Quality control prior to the dHCP neonatal fMRI pipeline

The following is adapted from the release notes for the second dHCP data release. For further information see: http://www.developingconnectome.org/release-notes/

QC on the released dHCP data is performed at numerous stages in the analysis, including within the dHCP neonatal fMRI pipeline as described in section 3.6. There are three QC stages implemented prior to the neonatal fMRI pipeline:

1. Scanning notes were recorded by the radiographers, and failed scans were manually flagged as pass/fail depending on if the issue affects the fMRI
2. After reconstruction the images were visually inspected and each image was flagged as PASS/FAIL
3. The structural pipeline QC combined several sources of information: 479 scans were scored visually as part of an atlas construction project - we excluded scans with more than minor motion artifacts in T2. We excluded 11 scans we knew to be in error. We excluded scans on which the structural pipeline failed to run, or on which the separate structural QC pipeline failed to run. We did a visual inspection of all white matter surfaces and excluded one scan that was obviously failing.

The dHCP-538 cohort used within this paper comprises 538 subjects that passed all three stages of QC prior to the fMRI pipeline.

### 9.3. Detailed registration methods

Primary registrations:

1. *fieldmap-to-structural:* rigidly align the derived fieldmap magnitude image (see Section 3.2) to the native structural T2w space using FSL FLIRT (Jenkinson et al., 2002b; Jenkinson and Smith, 2001). A boundary-based registration (BBR) (Greve and Fischl, 2009) cost function is used if the fieldmap was derived from the spin-echo EPI using TOPUP. However, the correlation ratio cost function is used if the fieldmap was derived from the gradient-echo, because the magnitude image lacked sufficient anatomical detail for BBR. The fieldmap-to-structural transform is then applied to resample the fieldmap image into the native structural space.
2. *sbref-to-structural:* rigidly align the single-band EPI image (sbref) with the native structural T2w space and correct for susceptibility distortions in the sbref using FSL FLIRT, with the BBR cost function, and FSL FUGUE. This step requires the fieldmap to be in the native structural space (calculated in the previous registration stage) to correct for susceptibility distortions in the sbref.
3. *functional-to-sbref (distorted):* rigidly align the functional multiband EPI image with the sbref using FSL FLIRT with the default correlation ratio cost function. This registration step is performed prior to susceptibility distortion correction of the functional multiband EPI as described in Section 3.4, therefore both the functional multiband EPI image and the sbref will contain susceptibility distortions. The first volume of the functional multiband EPI is used as the source (moving) image in this registration because the subsequent motion correction and distortion correction stage defines the functional space from the first volume (see Section 3.4).
4. *functional-to-sbref (undistorted)*: after motion correction and distortion correction, rigidly align the distortion-corrected functional multiband EPI image with the distortion-corrected sbref using FSL FLIRT with the default correlation ratio cost function. All volumes in the corrected functional multiband EPI image are aligned as consequence of the motion correction, therefore the temporal mean is used as the source (moving) image in this registration as it typically has superior SNR compared to a single volume.
5. *template-to-structural*: align the structural image to the dHCP volumetric template (Schuh et al., 2018b). Template-to-structural registration is performed with a multimodal non-linear registration (ANTs SyN)(Avants et al., 2008) of the age-matched T1w and T2w template to the subject’s T1w and T2w structural, which is then combined with the appropriate atlas week-to-week transforms to yield a (40 week) template-to-structural transform. We also evaluated FSL FNIRT (Jenkinson et al., 2012) and MIRTK Register (Similarity+Affine+FFD transformation model) (Schuh et al., 2018a) and found that Register achieved excellent alignment but was not sufficiently regularised, resulting in inversion inaccuracy, whilst FNIRT was well regularised but did not produce alignments with sufficient accuracy (data not shown). We expect that good results could be achieved with both tools if their parameters were optimised, however ANTs SyN provided a good trade-off between alignment and regularisation with minimal parameterisation. In the event that the age of the subject is outside the range covered by the atlas, the closest template age within the atlas is used. Furthermore, some subjects do not have a T1w image, so in this instance only the T2w is used.

Composite registrations:

1. *fieldmap-to-functional:* constructed by combining the fieldmap-to-structural transform with the inverse sbref-to-structural and inverse functional-to-sbref (distorted) transforms. This allows for the fieldmap to be resampled to the native functional space, which is essential for the subsequent motion correction and distortion correction (Section 3.4). We have found that aligning the fieldmap with the functional via the structural is very robust and precise, largely because both sub-steps use BBR cost functions.
2. *functional-to-structural (undistorted):* constructed by combining the functional-to-sbref (undistorted) affine with the sbref-to-structural affine, which yields a linear transform that aligns the motion and distortion corrected functional multiband EPI with structural T2w.
3. *functional-to-template (undistorted):* constructed by combining the functional-to-structural (undistorted) transform with the inverse template-to-structural transform to yield the functional-to-template (undistorted) non-linear transform to align the motion and distortion corrected functional multiband EPI with the 40-week dHCP template space with a single resampling.

### 9.4. Frame censoring

A popular and effective method of dealing with head motion is using spike regression (Satterthwaite et al., 2012) or scrubbing (Power et al., 2014, 2012), collectively referred to as frame censoring. We evaluated frame censoring as an alternative to ICA+FIX denoising.

Both spike regression and scrubbing first identify time-points (whole volumes referred to as frames) and then censor these frames so that they do not affect downstream analysis. The methods differ in how they identify the contaminated frames and how they censor the contaminated frames (Parkes et al., 2018). Both methods use framewise displacement (FD; see Table 3) with a fixed displacement threshold to identify contaminated frames. Scrubbing additionally uses DVARS (see Table 3), also with a fixed threshold. Censoring in spike regression is achieved by creating a nuisance regressor per contaminated frame, whereas scrubbing either excludes contaminated frames and/or replaces contaminated frames with surrogate data depending upon what is appropriate for downstream processing. Additionally, both techniques employ a heuristic that discards entire subjects if there are insufficient uncontaminated time-points.

Frame censoring methods can be expensive in terms of the number of volumes censored, particularly in high-motion cohorts such as neonates. This is particularly true for the dHCP because the babies are scanned without sedation. Using framewise displacement (Power et al., 2012) as a surrogate for head motion and a threshold of 0.25 mm, as advocated by Satterthwaite et al. (2013), results in ~20% of frames being flagged as motion corrupted. Furthermore, if we exclude subjects with < 4 minutes of continuous uncorrupted data, the minimum recommended in spike regression and scrubbing (Parkes et al., 2018; Power et al., 2014; Satterthwaite et al., 2013), then only 18 of 538 subjects are retained. Thus, to implement frame censoring we needed to relax these criteria. To identify contaminated frames we used DVARS (post motion and distortion correction), with a threshold defined as the 75th percentile + 1.5 times the inter-quartile range (the outlier whisker when creating boxplots). This resulted in 6.5% of frames being flagged as outliers. We did not implement a minimum duration of continuous uncorrupted data. To achieve censoring, we simply removed the contaminated volumes, because the downstream analysis plan was to perform group sICA which is not sensitive the discontinuous time.

Frame-censoring was evaluated on the 512 subjects from that dHCP-538 data that passed QC (see Section 3.6). Frame censored data was compared to the ICA+FIX denoised data (described in Section 3.5) using spatial and network matrix similarity to the unbiased group template (also described in Section 3.5). This required running a low-dimension group ICA (dimension=25) across all data (all subjects, frame-censored and ICA+FIX denoised) to generate unbiased group spatial maps. The unbiased group maps were visually inspected and 12 RSN consistent maps identified (see Supplementary Figure 4).

**Supplementary Figure 4.**
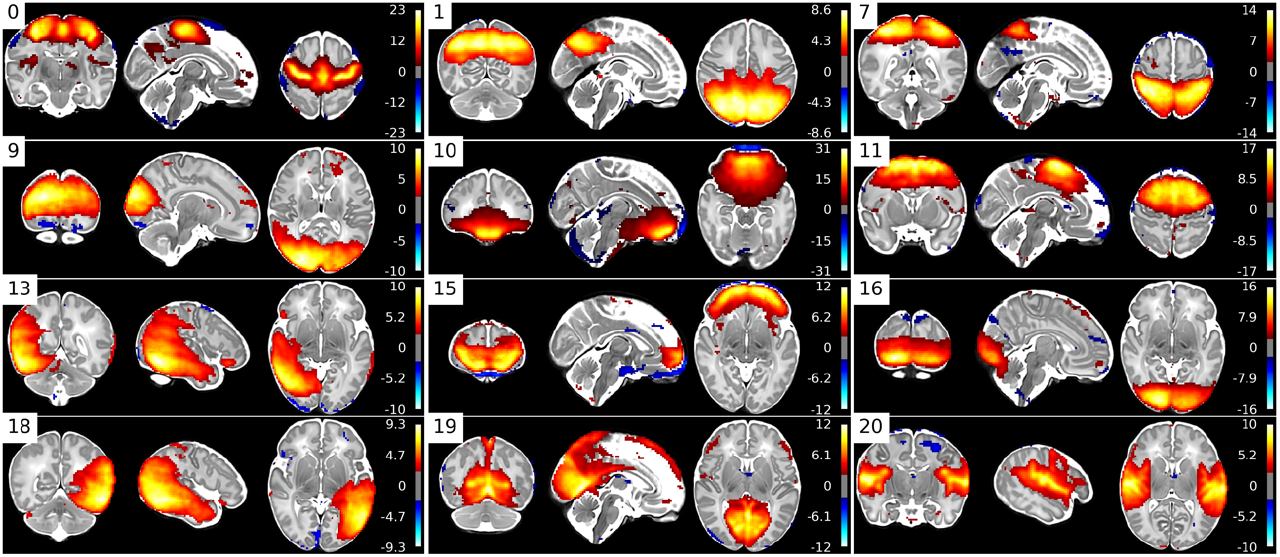
Unbiased group RSN template maps created from ICA+FIX and frame-censored data.

Group paired-differences in spatial and network similarity between frame-censored and ICA+FIX denoised groups were calculated using FSL RANDOMISE (Winkler et al., 2014) with 5000 permutations. Multiple comparison correction was achieved by FDR correction. ICA+FIX denoised data resulted in significantly (p<0.025) greater spatial similarity to 10 of the 12 unbiased group RSN maps, whilst frame censoring only showed significantly (p<0.025) greater spatial similarity for one of the unbiased group RSN maps (see Supplementary Figure 5). Furthermore, ICA+FIX denoising resulted in significantly (p<0.025) greater network matrix similarity to the unbiased group network matrix than frame censoring (see Supplementary Figure 6).

**Supplementary Figure 5.**
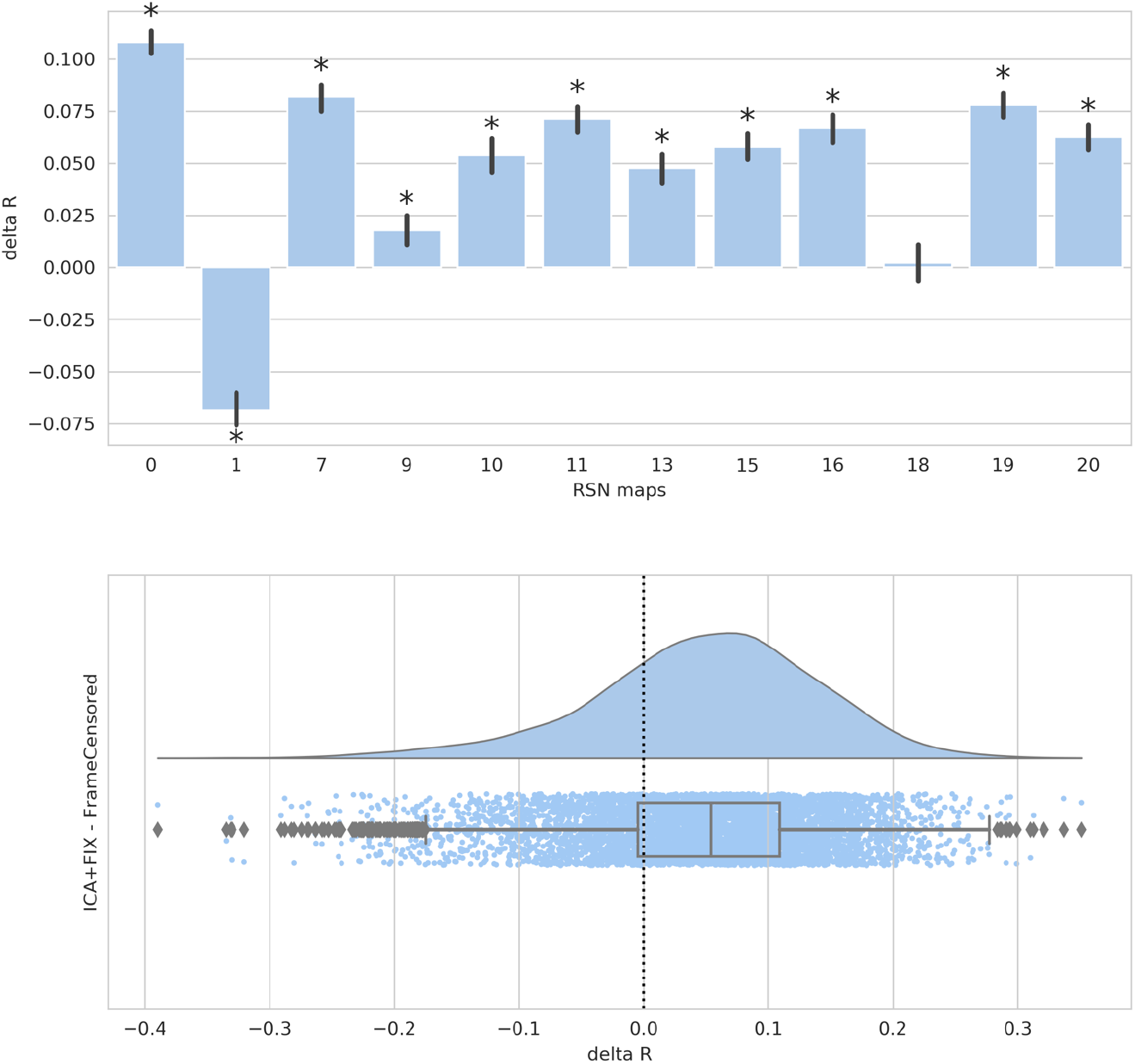
(Upper) Mean paired-difference (ICA+FIX minus Frame Censored) of spatial similarity to the unbiased group template per map. Asterisks indicate significant differences. (Lower) Distribution of paired differences (ICA+FIX minus Frame Censored) of spatial similarity to the unbiased group template pooled over all spatial maps.

**Supplementary Figure 6.**
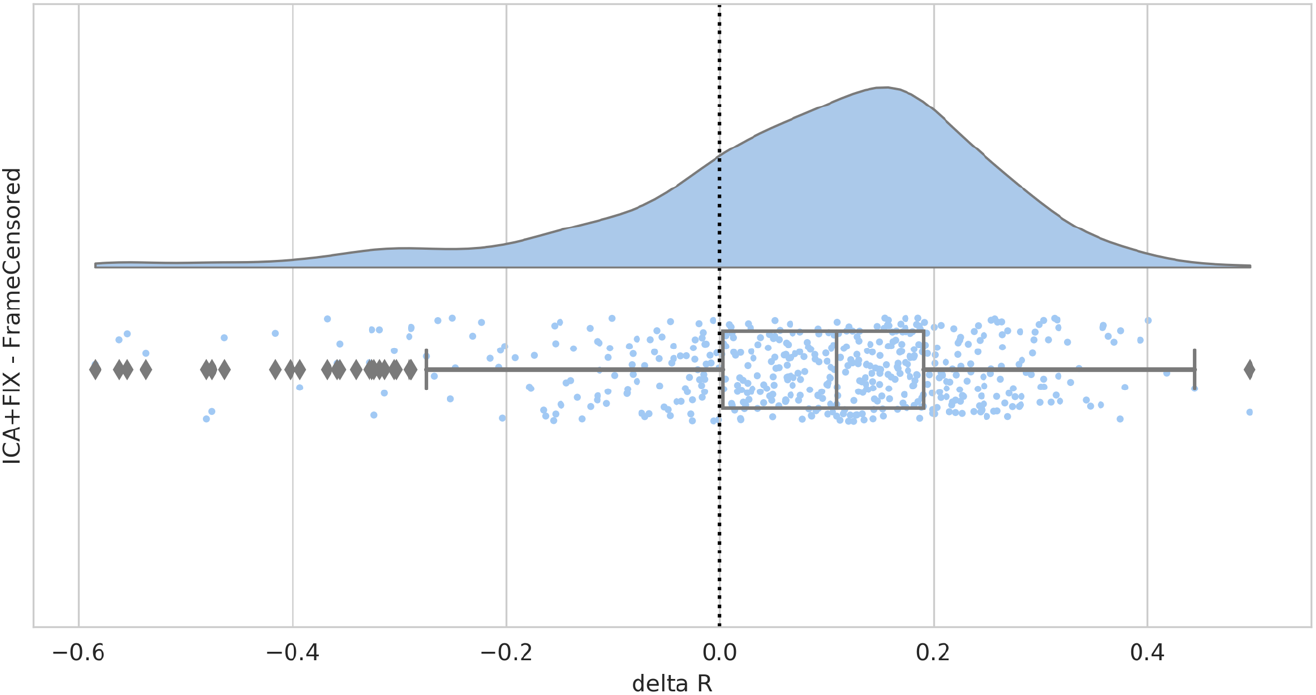
Distribution of paired differences (ICA+FIX minus Frame Censored) of network matrix similarity to the unbiased group network matrix.

These results suggest, that in this context, with this specific variant of frame censoring, that ICA+FIX denoising performs better than frame censoring on these specific benchmarks. However, ICA+FIX has the added advantage that it is able correct for confounds beyond motion, such as multi-band artefacts, scanner artefacts, venous/arterial related artefacts, and spin-history effects. Thus, given this evidence, we have opted to implement ICA+FIX in this dHCP neonatal fMRI pipeline which enables us to largely mitigate the effects of many confounds without excluding any subjects or time-points. Furthermore, we have avoided introducing a hard-censoring step at an intermediate processing point, which could have ramifications for downstream processing. It is important to note that this analysis was intended to provide readers with a sense for how the results from the two techniques compare, but does not necessarily provide definitive results regarding the performance of the two approaches in all situations, with all possible settings and analysis decisions included, as we only consider a single robust implementation of frame censoring and a specific set of outcome metrics. We expect both approaches could be valuable and effective for infant fMRI processing in the right context, and both likely better than doing neither.

### 9.5. RSN development with age

Before regressing the RSNs on age to look for developmental changes, we examined a number of age-related confounds. Specifically, we correlated DVARS and FD (as surrogates for motion), tSNR, and brain volume (estimated as the number of voxels in the subject’s brain mask in func space). We observed a strong positive correlation of brain volume with age (r=0.86), a small positive correlation of mean DVARS (r=0.05) and mean FD (r=0.16) with age, and a small negative correlation of tSNR (r=-0.15) with age (see Supplementary Figure 7). Movement and tSNR are clear confounds that we wish to control for, however brain volume is more challenging because it can be both a confound (due to differences in relative resolution and signal) and a legitimate feature of development. Here we control for brain volume and present RSN correlations with development.

**Supplementary Figure 7.**
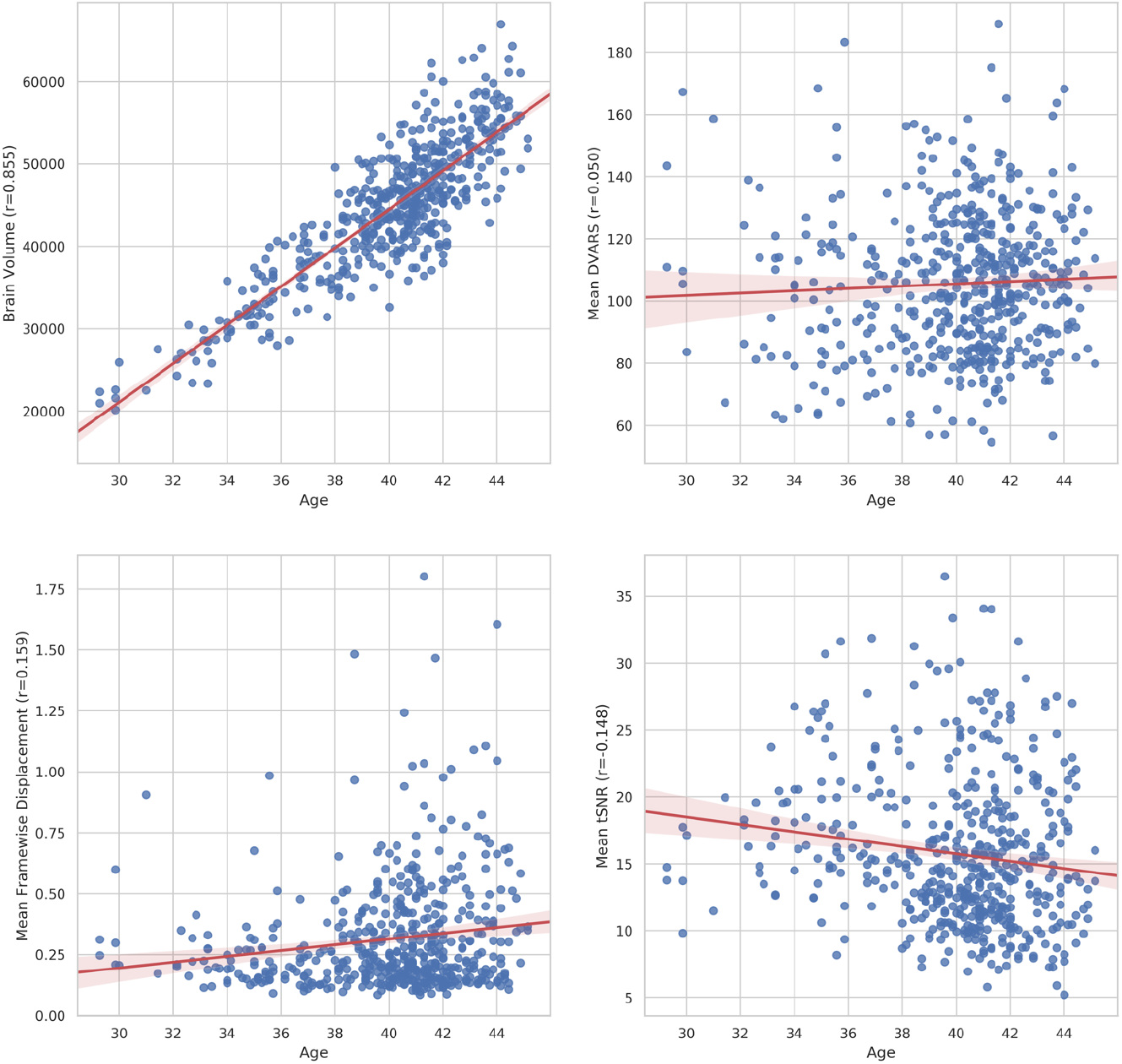
Age-related confounds. Age is the post-menstrual age-at-scan in weeks.

We used FSL dual-regression (DR)(Nickerson et al., 2017) to regress all the PFM-group-maps onto the individual subject fMRI to yield subject-specific time-courses and spatial maps. As recommended by Nickerson et al. (2017), when performing DR the subject-specific time-courses were variance normalised before the second stage of DR which means that the single-subject DR spatial maps capture both the spatial distribution of the network (i.e. “shape”) as well as the “amplitude” of the network activity. To allow us to delineate the contribution of just amplitude alone, we additionally calculated the median absolute deviation of the DR time-courses (not variance-normalised) as an estimate of amplitude. To investigate changes with age, we regressed the spatial maps and amplitudes on age, controlling for DVARS, FD, tSNR, and brain volume (see Supplementary Figure 8) using FSL RANDOMISE (Winkler et al., 2014) with 5000 permutations. Multiple comparison correction was achieved by FDR correction with a threshold of 0.05.

The DR spatial maps show a significant effect for age in all modes, in voxels that are spatially consistent with the group PFM map. Furthermore, the DR amplitudes show a significant increase in network amplitudes with age for all modes, which indicates that the age effects are, at least partially, driven by this increased amplitude of network activity.

**Supplementary Figure 8.**
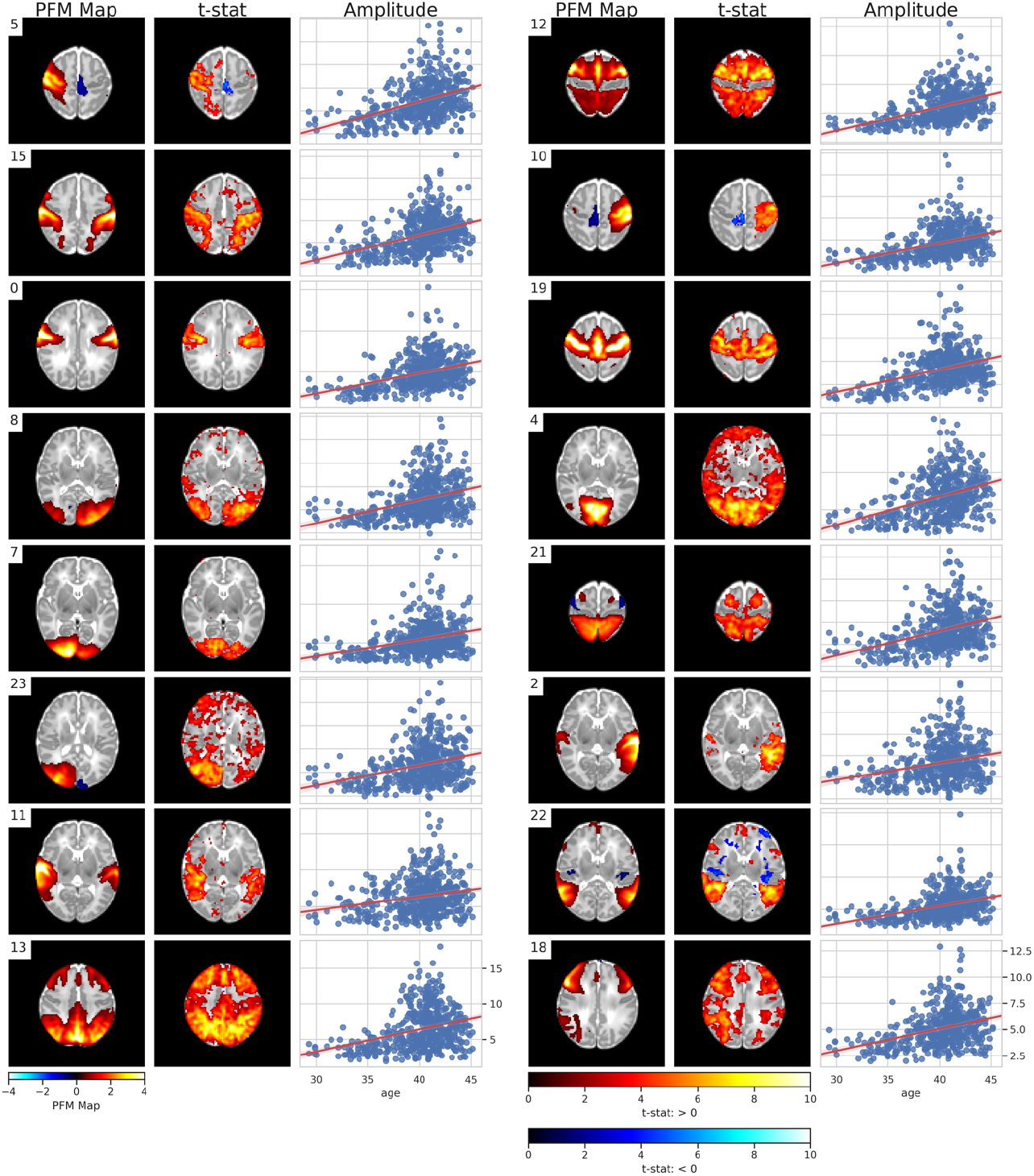
PROFUMO group spatial maps (PFM Map), t-statistic of age regressed on the DR spatial maps (t-stat), and DR amplitudes with age (Amplitude) for the 16 modes qualitative assessed as corresponding to adult resting-state networks. Age is the post-menstrual age-at-scan in weeks. Brain volume, mean DVARS, mean tSNR, and mean FD confounds are controlled. Only significant results are shown. Multiple comparison correction was achieved by FDR correction with a 0.05.

## 10. Supplementary Figures

**Supplementary Figure 9.**
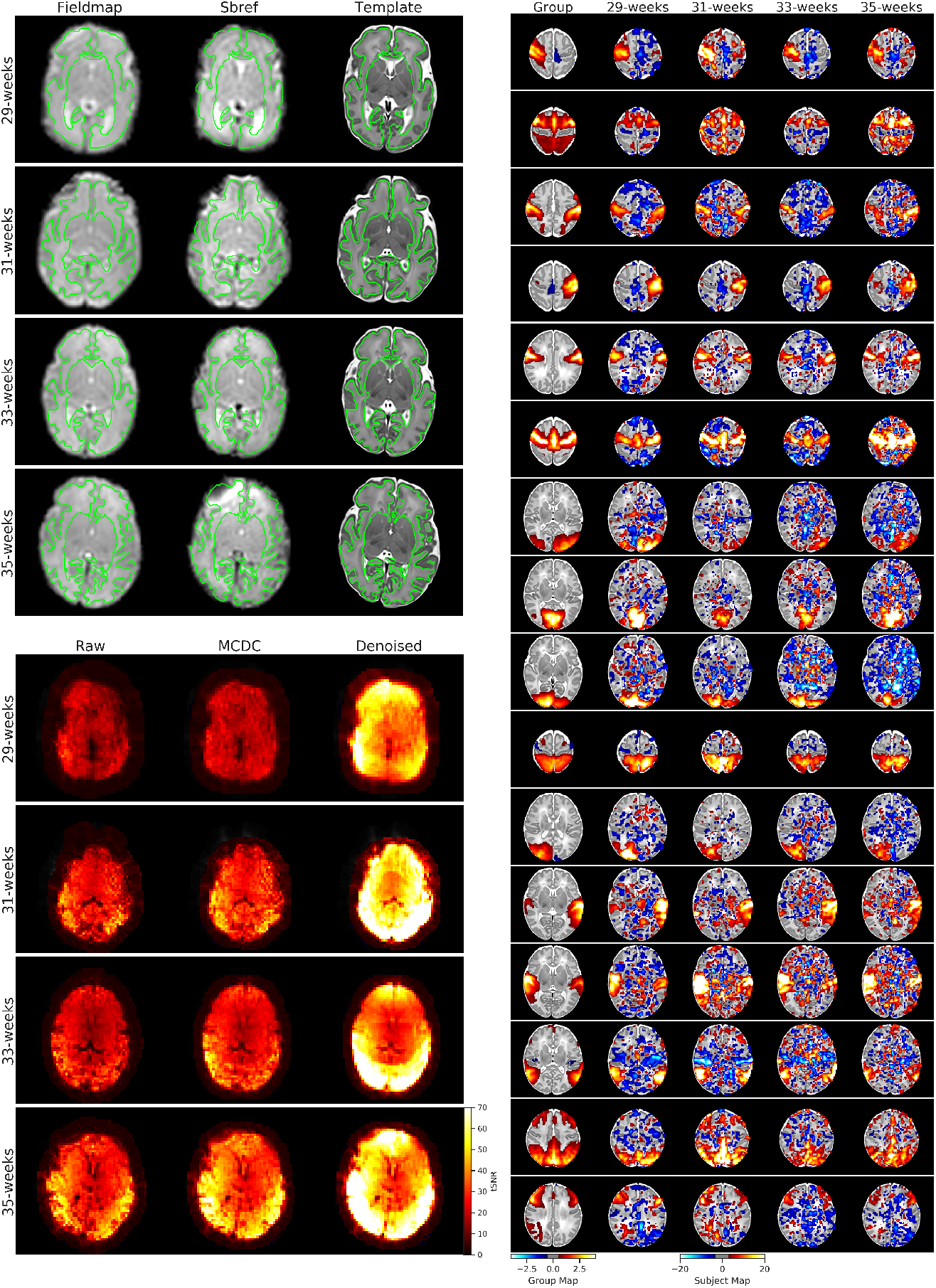
Examples of pre-term subjects scanned at 29-weeks (born at 28-weeks), 31-weeks (born at 30-weeks), 33-weeks (born at 31-weeks), and 35-weeks (born at 34-weeks). Upper left images demonstrate registration quality for these subjects. Fieldmap, sbref, and template images are resampled to the native structural reference space. The outline of the native structural white matter discrete segmentation is overlaid in green. Lower left images demonstrate mean tSNR for these pre-term subjects at different pipeline stages: raw EPI, motion and distortion corrected EPI (MCDC), and denoised EPI. The right images demonstrate the single-subject spatial maps for these subjects after dual-regressing 16 profumo group RSN maps onto them. The first column is the group map, and the subsequent columns are the dual-regressed single subject maps.

**Supplementary Figure 10.**
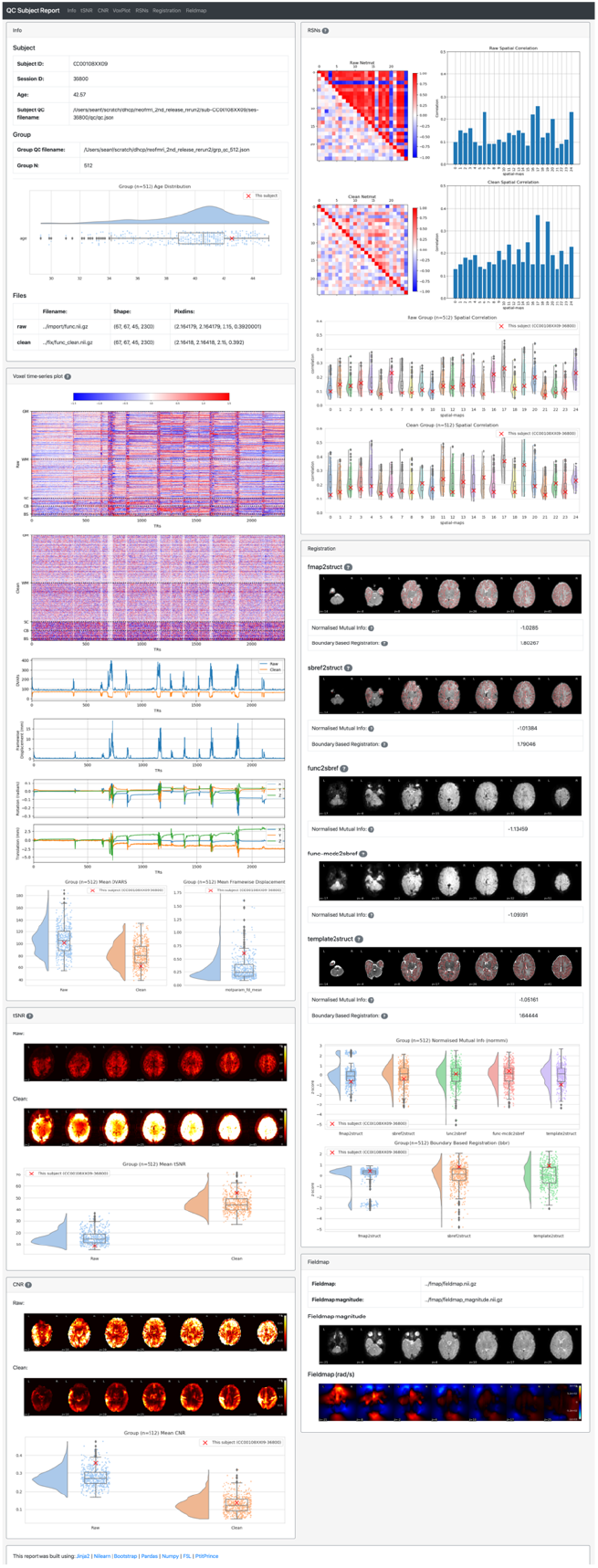
Screen shot of automated QC report for a single subject. The dHCP neonatal fMRI pipeline automatically calculates a number of QC metrics (MP, DVARS, FD, tSNR, CNR, NMI; see Table 3) and generates this HTML QC report for each subject. The report presents each QC metric for the individual within the context of the group distribution for the corresponding metric. Additionally, the report also presents descriptive/qualitative summaries of the subject’s data quality in the form of “voxplots “ and spatial maps for each metric as applicable. The report generation tool utilises a variety of open source packages including Jinja2 (http://jinja.pocoo.org/docs/2.10/), Bootstrap (https://getbootstrap.com/), Pandas (https://pandas.pydata.org/), Numpy (https://www.numpy.org/), Seaborn (https://seaborn.pydata.org), Nilearn (http://nilearn.github.io/), FSL (https://fsl.fmrib.ox.ac.uk/fsl/fslwiki), and PtitPrince (https://github.com/pog87/PtitPrince).

**Supplementary Figure 11.**
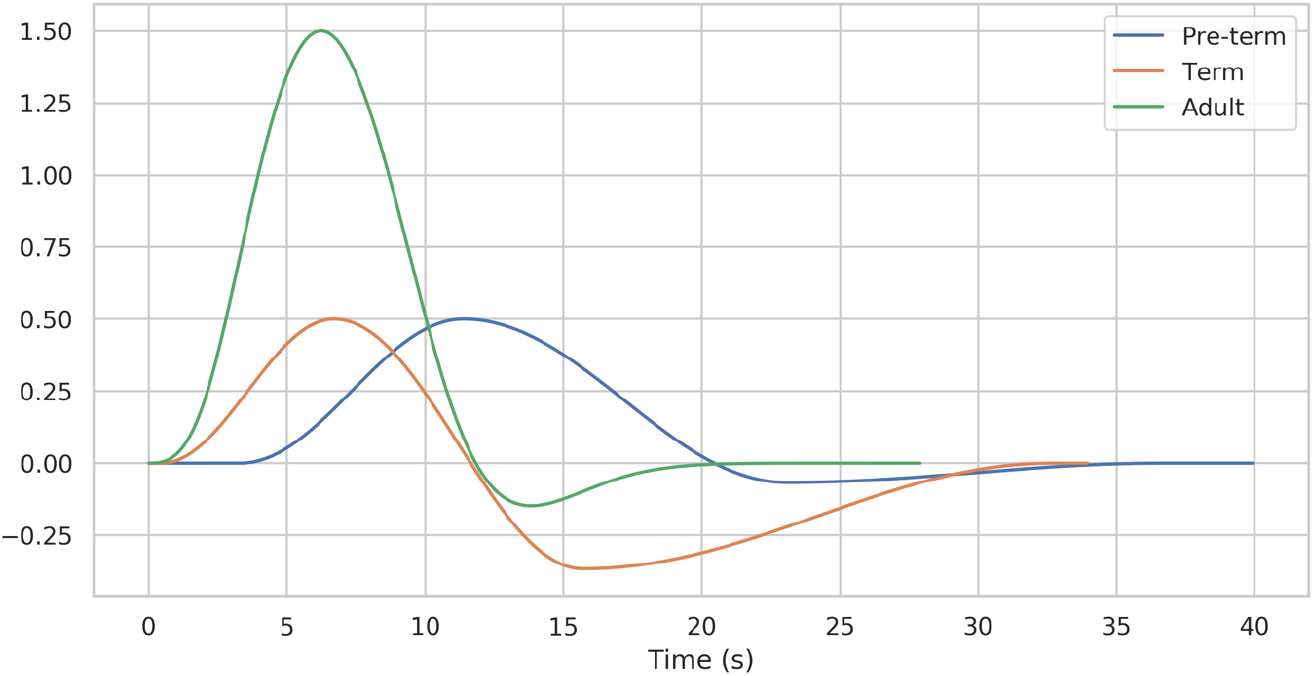
Term and pre-term HRF models constructed for this study, and the default adult HRF model within PROFUMO. The term and pre-term haemodynamic response characteristics are adapted from Arichi et al. (2012). The amplitude of each HRF is rescaled as part of the fitting process, however, for visualisation purposes the peak amplitudes here have been scaled to be consistent with the measured amplitudes in Arichi et al. (2012).

